# HLH-30/TFEB is necessary for chromatin reorganization and maintenance of cell quiescence during starvation in *C. elegans*

**DOI:** 10.1101/2025.10.31.685810

**Authors:** Marta Muñoz-Barrera, Alejandro Mata-Cabana, Almudena Moreno-Rivero, Francine A. Piubeli, Beatriz Ren-Barroso, Nada Al-Refaie, Gabriel Gutierrez, Daphne S. Cabianca, María Olmedo

**Author notes:** These authors contributed equally (Marta Muñoz-Barrera, Alejandro Mata-Cabana).

## Abstract

Cellular quiescence is a metabolically active, non-proliferative state critical for tissue maintenance and regenerative capacity, with broad implications for aging and age-related diseases. In *Caenorhabditis elegans*, L1 developmental arrest upon hatching in the absence of food provides a robust in vivo model to study quiescence. Here, we investigate the roles of the transcription factors HLH-30/TFEB and DAF-16/FOXO during L1 arrest. We show that HLH-30 and DAF-16 collaborate to ensure survival under starvation, with reciprocal regulation of their subcellular localization and transcriptional activity. HLH-30 exerts broad transcriptional control during L1 arrest, modulating genes involved in chromosome organization and cell cycle progression. Profiling of chromatin spatial distribution reveals that HLH-30 is required for fasting-induced 3D chromatin reorganization. Loss of HLH-30 disrupts seam cell cycle arrest and leads to overactivation of the pioneer transcription factor BLMP-1, leading to premature initiation of developmental programs under starvation. Our findings uncover previously unrecognized functions of HLH-30 in genome architecture and quiescence regulation, highlighting conserved mechanisms of transcriptional control during nutrient deprivation with implications for aging and disease.

## INTRODUCTION

Cell quiescence is a reversible non-proliferative state maintained by cell cycle inhibitors. Despite the lack of proliferation, quiescence is a metabolically active process involving protective mechanisms to ensure survival and enable re-entry into the cell cycle (reviewed in ^1^). This process is crucial for various cell types, particularly for preserving adult stem cell pools, thereby contributing to tissue homeostasis and regeneration ^2^. Adult stem cells can reside in tissues for decades, but their survival and reactivation capacity diminish over time ^3,4^. Aged stem cells accumulate reactive oxygen species (ROS), DNA damage, protein aggregates, and show mitochondrial malfunction ^5^. These cellular damages impair the reactivation of quiescent cells, reducing regenerative capacity as we age ^6–8^.

To study cell quiescence in vivo, we use the developmental arrest of *C. elegans* L1 larvae as a model ^6,9,10^. During *C. elegans* late embryogenesis, stem-like cells arrest prior to hatching ^11^. When embryos hatch in the absence of food, development pauses at the L1 stage. These cells remain arrested until nutrients become available to allow post-embryonic development to commence. During L1 arrest, larvae exhibit aging markers such as increased ROS, protein aggregation, and mitochondrial fragmentation ^6^. After prolonged L1 arrest, aged quiescent larvae experience delays in restarting proliferation due to a latency phase in the division of stem cell-like blast cells (seam cells) ^12^. This process parallels cellular quiescence in higher eukaryotes ^9^. Fasting at the L1 stage also provokes a large-scale chromatin reorganization ^13^.

The transcription factor DAF-16, the *C. elegans* ortholog of mammalian FOXO3A (Forkhead box O3), plays a crucial role during L1 arrest of *C. elegans*. DAF-16 acts at the beginning of L1 arrest by activating stress response pathways (reviewed in ^14^). Its loss compromises survival and recovery from L1 starvation ^15^. Another transcription factor, HLH- 30, the ortholog of mammalian TFEB (Transcription Factor EB), is also involved in metabolic regulation during L1 starvation ^16,17^. Interestingly, depending on the physiological context, DAF-16/FOXO and HLH-30/TFEB function synergistically or have opposing effects ^18^, but their combined effect had never been tested during L1 arrest.

We investigated the roles of the transcription factors (TFs) DAF-16 and HLH-30 during L1 arrest by analyzing single and double mutants. Our findings reveal a reciprocal influence between these factors, demonstrated through changes in subcellular localization and global gene expression patterns. Notably, we uncovered previously unknown functions of HLH-30 in regulating L1 arrest. HLH-30 modulates gene expression linked to chromosome organization and cell cycle progression. An analysis of spatial chromatin distribution further showed that HLH-30 is essential for fasting-induced chromatin reorganization. Moreover, *hlh-30* mutants exhibited impaired seam cell cycle arrest under starvation, a defect partially attributed to the overactivation of the pioneer transcription factor BLMP-1. Overall, our study highlights novel roles for HLH-30 in maintaining cell quiescence and shaping genome architecture. This research sheds light on conserved mechanisms of transcriptional regulation during cellular quiescence and nutrient deprivation—processes that are fundamental to aging, metabolic disorders, and cancer.

## RESULTS

### HLH-30 and DAF-16 collaborate to allow survival to L1 arrest

To investigate a possible collaborative role of DAF-16 and HLH-30 during L1 starvation (Fig. 1A), we first analyzed the subcellular localization of DAF-16 and HLH-30 using the fluorescent reporters *ot971[daf-16::gfp] and sqIs17[hlh-30p::hlh-30::gfp]* (Fig. S1A). As previously shown with a DAF-16 multicopy reporter ^19,20^, DAF-16::GFP showed high levels of nuclear localization at the beginning of L1 starvation, and these levels decreased over time in starvation (Fig. 1B). In *hlh-30* mutants, DAF-16 nuclear localization was significantly reduced when compared to the wild-type background at day 1 and 4 (Fig. 1B). Localization at later stages of arrest is not assessable due to the limited survival of the mutants. In contrast, HLH-30::GFP exhibited approximately 50% nuclear localization on day 1 of L1 arrest, which declined by day 4 before increasing again at later stages of arrest (Fig. 1C). In the *daf-16* mutant background, HLH-30 showed higher nuclear localization than in the wild type (Fig. 1C). These data show that DAF- 16 and HLH-30 subcellular localization depend on each other and suggest that a collaboration between both TFs could occur at the beginning L1 starvation, when both TFs are found in the nucleus. Indeed, previous analysis of DAF-16 and HLH-30 direct and indirect targets over L1 arrest revealed that the transcriptional response controlled by this transcription factors is mounted early during L1 starvation ^21^.

**Figure 1.**
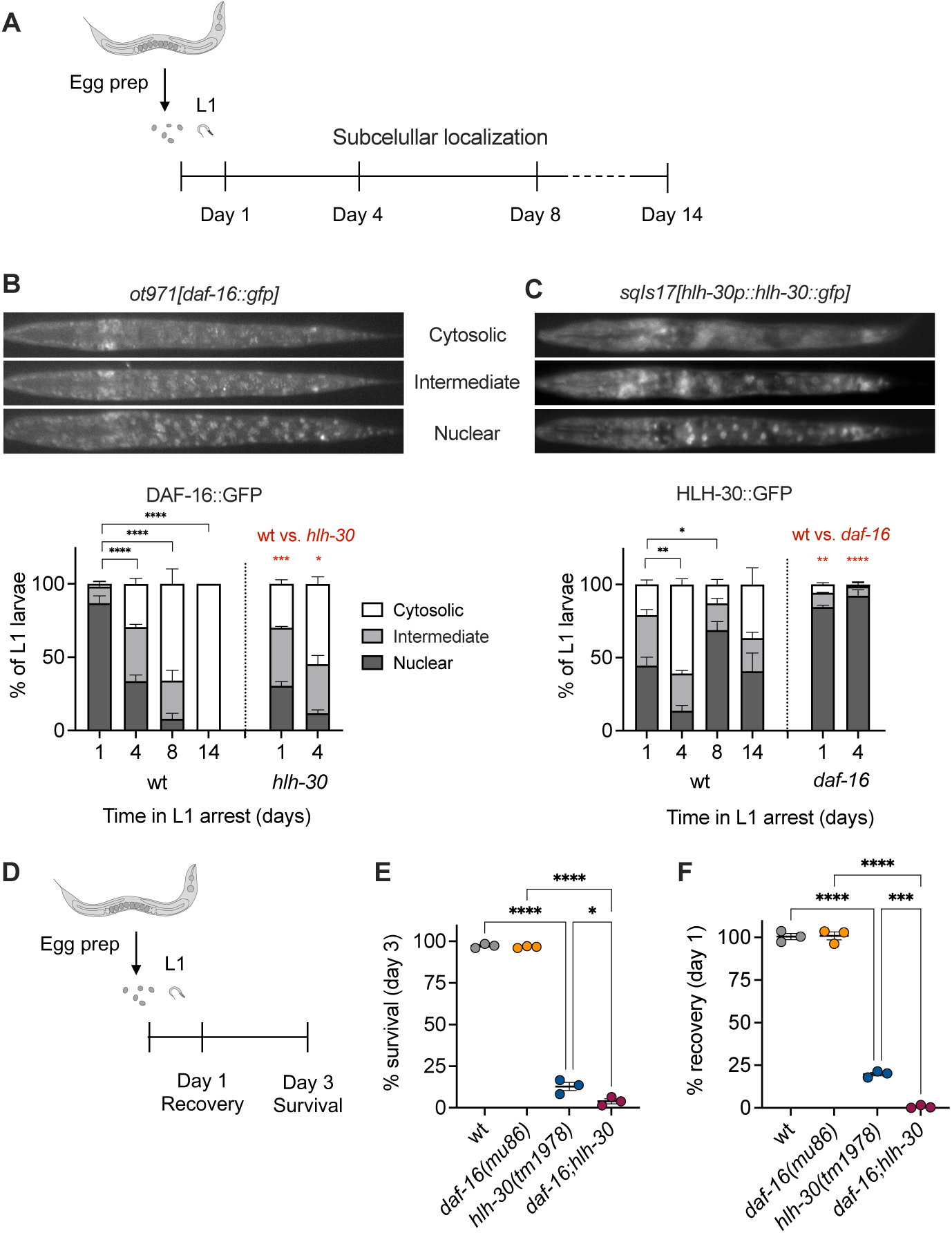
DAF-16 and HLH-30 play a role in the first hours of starvation. **(A)** Time points for the analysis of the subcellular localization of DAF-16::GFP and HLH-30::GFP. **(B-C)** Representative images and quantification of the subcellular localization at different times of L1 arrest of DAF-16::GFP (B) and HLH-30::GFP (C). Each larva in a population was categorized as showing cytosolic, intermediate or nuclear localization of the corresponding TF. Error bars represent SEM. We first compared the averages of nuclear localization by one-way ANOVA followed by Dunnett’s multiple comparisons to test the differences between day 1 and longer times of arrest in the wild type-background (marked in black). We used *t-test* to compare day 1 and 4 of the mutant samples to the corresponding days of the wild type (red symbols) **(D)** Time points for analysis of recovery and survival of the wild type, the *daf-16(mu86)* and *hlh- 30(tm1978)* mutant, and the corresponding double mutant. **(E)** Percentage of L1 larvae alive after 3 days of starvation for the wild-type, the *daf-16(mu86)* and *hlh-30(tm1978)* mutant, and the corresponding double mutant. **(F)** Percentage of L1 larvae that resume development upon refeeding after 1 day of starvation, for the wild-type, the *daf-16(mu86)* and *hlh-30(tm1978)* mutant, and the corresponding double mutant. Colored dots represent results from independent experiments, and the lines mark the mean (±SEM). In all cases, we performed One-way ANOVA followed by Tukeýs multiple comparisons to compare all samples to each other. Only significative differences are shown.

We then evaluated the impact of DAF-16 and HLH-30 on survival and in the capacity to recover from L1 starvation. Since the mutant *hlh-30(tm1978)* shows impaired survival and recovery even after short periods of arrest ^16^, we focused on measuring survival after 3 days and recovery after 1 day of arrest (Fig. 1D). As expected, after this short period of starvation, the *daf-16(mu86)* was indistinguishable from the wild type, but *hlh-30(tm1978)* mutants showed strong defects in survival and recovery. Despite the apparent dispensability of DAF-16 to survive short arrest, the analysis of the double mutant showed that the *daf-16(mu86)* mutation aggravated the defects of *hlh-30(tm1978)* (Fig. 1E-F). These results suggests that there might be some degree of redundancy between DAF-16 and HLH-30 at the beginning of L1 arrest. Indeed, at short times of L1 arrest, activation of DAF-16 with a *daf-2* mutation (Fig. S1B) improved survival and recovery of the *hlh-30* in a DAF-16 dependent manner (Fig. S1C), suggesting that DAF-16 can take over some HLH-30 functions.

### HLH-30 exerts pervasive transcriptional control at L1 arrest

To understand the roles of DAF-16 and HLH-30 at the initial stages of L1 arrest, we performed mRNA sequencing of the wild type, *daf-16(mu86)*, *hlh-30(tm1978)* and the *daf-16;hlh-30* double mutant at day 1 of starvation. The initial evaluation of expression dynamics by principal-component analysis (PCA) showed that the biological replicates clustered together. Furthermore, the *daf-16* mutant is positioned close to the wild-type strain along PC1, in contrast to *hlh-30* and the double mutant (Fig. 2A). This suggests that the effect of the *hlh-30* mutation on the double mutant is stronger than that of the *daf-16* mutation. Indeed, we found a greater transcriptional deregulation in the *hlh-30* and double mutants than in the *daf-16* mutant strain. Specifically, in comparison to the wild type, we found 2975 differentially regulated genes in the *daf-16* mutant, 7899 genes in the *hlh-30* mutant and 10188 genes in the double *daf-16;hlh-*30 mutant (Fig. 2B, Fig. S2A-C, File S1). Most of the deregulated genes in the single mutants were also found in the double mutant (72% for the *daf-16* mutant, and 90 % for the *hlh-30* mutant). The high level of gene deregulation and the strong overlap between the *hlh-30* and the double mutants suggest a high similarity between these two conditions (Fig. 2B).

**Figure 2.**
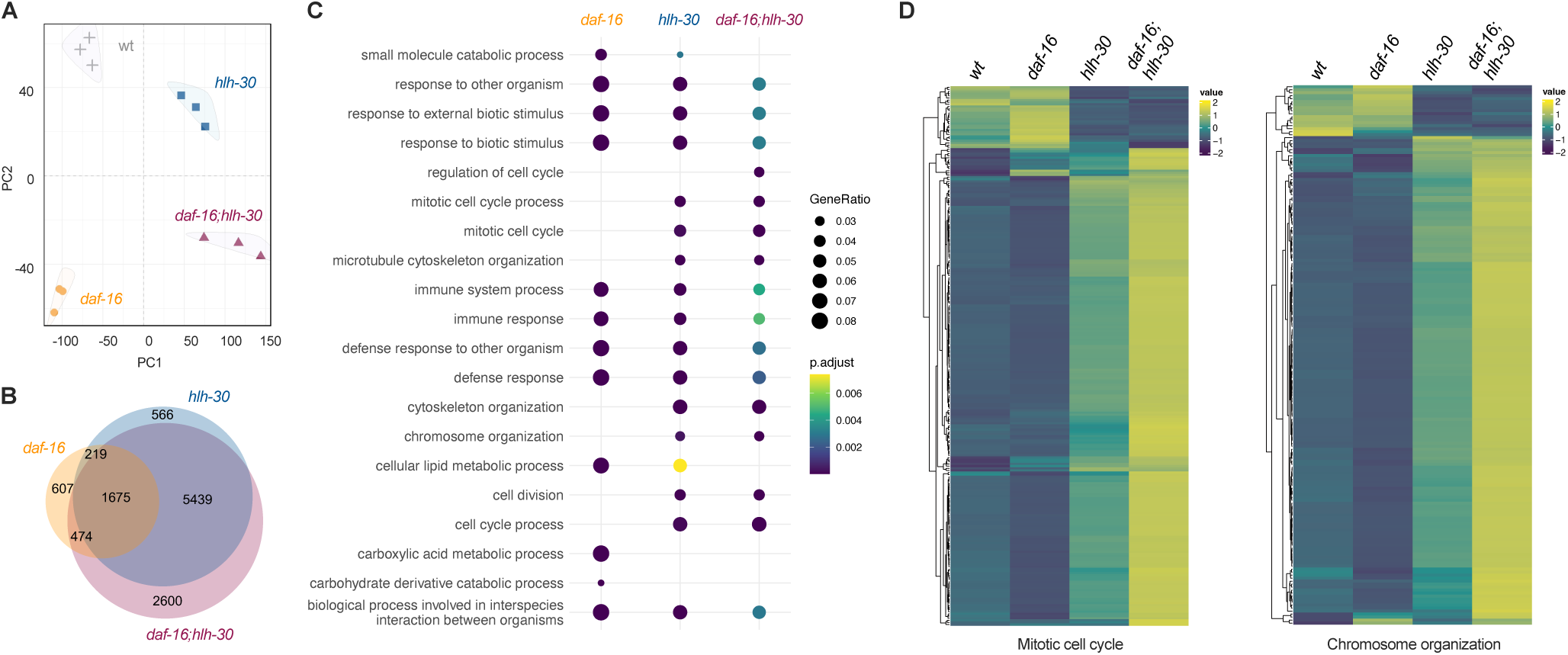
DAF-16 and HLH-30 impact transcription during L1 starvation. **(A)** Principal- component analysis of mRNA-seq data for all the four strains and three independent biological replicates. **(B)** Venn diagram showing the number of differentially expressed genes for each of the mutants. **(C)** Representation of the top 20 most significant biological processes affected across all samples. **(D)** Heatmaps showing the expression of genes within the categories “Mitotic cell cycle” (256 genes) and “Chromosome organization” (185 genes) in the double *daf-16;hlh-30* mutant sample.

Among the differentially expressed genes (DEGs), we observed that well-known specific DAF-16 and HLH-30 targets were deregulated exclusively in the samples corresponding to the *daf-16* and *hlh-30* single mutants respectively (Fig. S2D-E). This observation validated our experimental approach. We also detected changes in gene expression that suggest different types of collaboration between DAF-16 and HLH-30. Some genes required both DAF- 16 and HLH-30 for full activation (Fig. S2F), pointing to a collaborative function of these transcription factors. In contrast, other genes showed reduce expression only in the double mutant (Fig. S2G), suggesting a redundant role between DAF-16 and HLH-30 at day 1 of L1 arrest.

To identify biological processes associated with transcriptional activation, we performed functional enrichment analysis on the gene lists of DEGs with padj<0.05, for each mutant strain. This was achieved through the implementation of the clusterProfiler package (v4.14.6) ^22^. The annotation source used was the Gene Ontology (GO) terms for Biological Processes. The top 20 most significant categories were selected for representation. Results for the *daf-16* mutant showed a significant enrichment for categories related to defense response, cellular catabolism, and lipid metabolism (Fig 2C and S3A). Strikingly, although HLH- 30 had been described as a master regulator of lysosome biogenesis and autophagy ^23^, the categories with greater representation for the *hlh-30* and the double *daf-16;hlh-30* mutants were related to two groups: cell division and defense response (Fig 2C, S3B-C). The categories grouped within defense response are shared by all the three mutants, what could reveal cooperation between these two transcription factors, as previously reported in a different context ^18^. However, the enrichment in cell division categories appears to be specific to the absence of HLH-30 (Fig. 2D), suggesting a role for this protein in the control of this biological process at the onset of L1 arrest. Among the biological processes affected specifically in the *hlh-30* and the double *daf-16;hlh-30* mutants was also chromosome organization (Fig. 2D, File S1). This suggests that HLH-30 might regulate chromatin on a large scale, what could explain the large impact of the *hlh-30* mutation on gene transcription (Fig. 2B).

### HLH-30 is necessary for fasting-induced chromatin reorganization

Interestingly, fasting at the L1 stage provokes a large-scale spatial reorganization of chromatin in *C. elegans* intestinal cells, that can be reversed upon nutrient supplementation. This spatial reorganization induced by fasting requires inhibition of the mTORC1 pathway. Indeed, *let- 363*/TOR RNAi treatment or depletion of DAF-15/Raptor lead to fasted-like chromatin organization in fed animals ^13^. However, the elements that control chromatin reorganization downstream of mTORC1 remain unknown. Since HLH-30 is activated upon mTOR inhibition, we investigated a possible role of HLH-30 on chromatin reorganization during L1 starvation.

We analyzed chromatin architecture in intestinal cells by performing live imaging of HIS-72/H3.3 fused to GFP, as previously done ^13^ for the wild type, *daf-16(mu86)*, *hlh- 30(tm1978)* and the double mutant *daf-16;hlh-30* larvae on day 1 of L1 arrest (Fig. 3A). In wild type animals, we observed the previously reported reorganization of the intestinal genome into two concentric chromatin rings ^13^ (Fig. 3A–C). In contrast, all mutant conditions displayed altered chromatin organization, albeit to varying degrees, with *daf-16* mutants deviating less from the fasting-induced wild type pattern than *hlh-30* mutants (Fig. 3A–C, Fig. S4A-C). Interestingly, the chromatin profile of the *daf-16;hlh-30* double mutant did not show an additive effect of the two mutations, as the profiles of *hlh-30* and *daf-16;hlh-30* were nearly identical, with a correlation coefficient of 0.9988 (Fig. S4A). Together, these results indicate that although both DAF-16 and HLH-30 contribute to fasting-induced chromatin reorganization, they likely act through overlapping mechanisms, with HLH-30 playing the more prominent role.

**Fig. 3.**
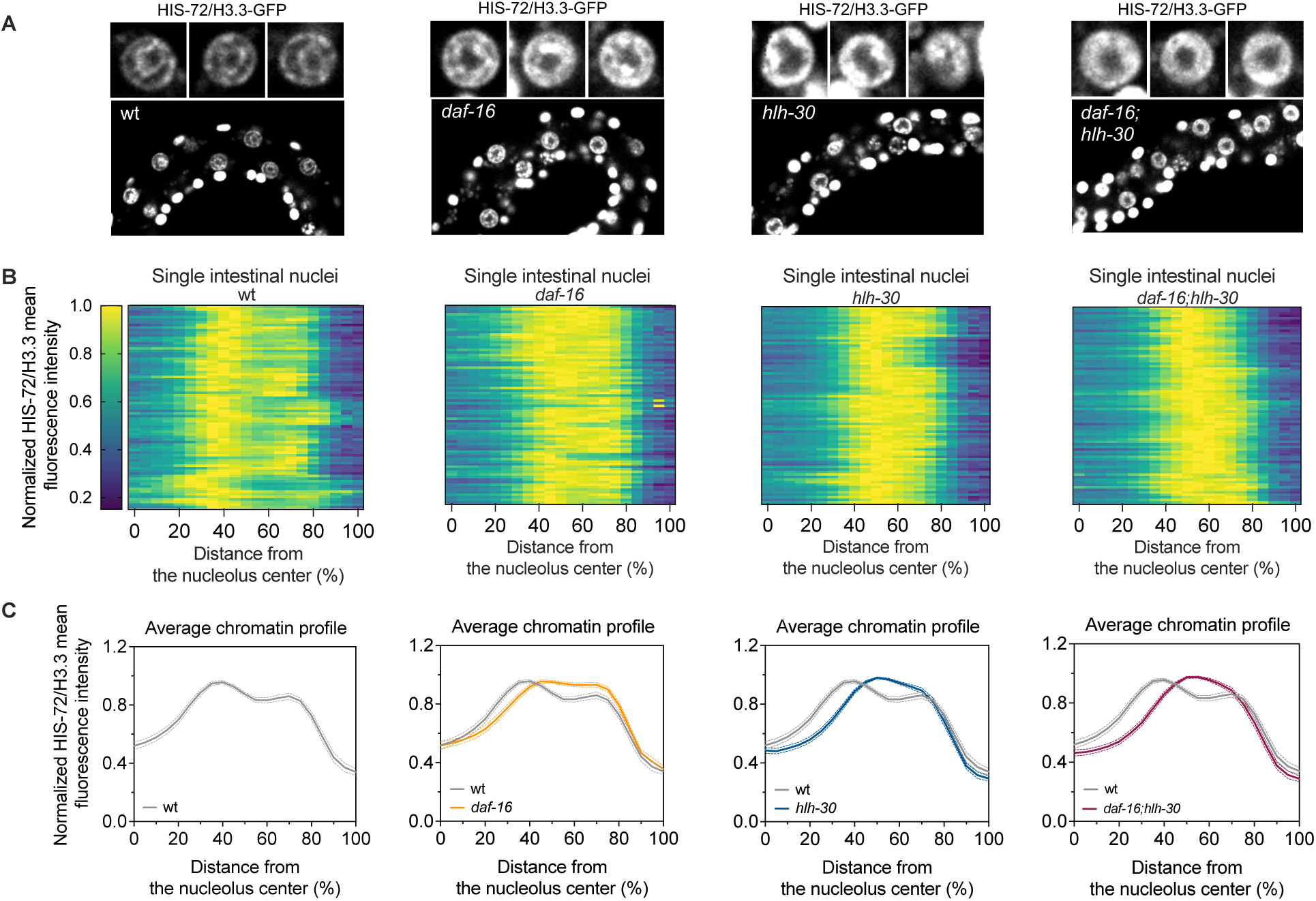
The *hlh-30(tm1978)* mutation impairs chromatin reorganization upon starvation. **(A)** Representative pictures of starved L1 larvae expressing the HIS-72/H3.3-GFP in a wild-type background, the *daf-16(mu86)* and *hlh-30(tm1978)* mutants, and the corresponding double mutant. Top: Representative intestinal nuclei. Bottom: Representative section of L1 larvae. **(B)** Heat maps showing HIS-72/H3.3–GFP fluorescence intensity in individual intestinal nuclei, with each row corresponding to one nucleus, as a function of the distance from the center of the nucleolus, which is centrally located in the intestine. Each plot includes a total of 78 nuclei from larvae tested in 3 independent experiments. **(C)** Averaged fluorescence intensity of the nuclei shown in (B). The dashed lines represent the 95% confidence interval of the mean profile. Average profiles were compared to estimate the statistical significance of differences between the mutants and the wild type, and the p*-*values are provided in Fig. S4C.

### HLH-30 is necessary to maintain cell quiescence during L1 arrest

Other interesting categories enriched in *hlh-30* and *daf-16;hlh-30* mutants were those related to cell cycle and cell division. This result was unexpected and particularly relevant given that the HLH-30 ortholog TFEB has diverse roles in cancer, functioning both as an oncogene and as a tumor suppressor, in lysosome-dependent and -independent manners ^24^. Furthermore, cell cycle arrest at the G1 phase is a relevant process during L1 starvation in *C. elegans*. The *hlh-30* mutant shows deregulation of multiple cell cycle-related genes (Fig. 2D), suggesting that HLH- 30 could play a role in the regulation of cell quiescence during L1 starvation. However, a role for HLH-30 in cell cycle quiescence had been discarded based on the analysis on M mesoblast cell divisions during L1 arrest ^16^.

Here, we decided to analyze the state of seam cells during starvation, as the *C. elegans* hypodermis conveys nutritional information to control cell quiescence ^25^. During postembryonic development of fed larvae, seam cells divide before the M mesoblast. We used a double reporter to follow the divisions of seam (P*scm::gfp*) and M cells (P*hlh-8::gfp*) in the wild type, and the *daf-16(mu86)*, *hlh-30(tm1978),* and double mutants. We calculated the fraction of starved larvae with divisions after one and two days of L1 starvation (Fig. 4A). For M cells, we did not observe any divisions at day 1 or day 2 in any of the strains (not shown). When analyzing seam cells, the *daf-16* mutant did not show a significant level of divisions compared to the wild type at day 1 or day 2. In contrast, after two days of starvation, the *hlh- 30* mutant showed a higher percentage of larvae with seam cell divisions compared to the wild type and the *daf-16* mutant. In the case of the *daf-16(mu86); hlh-30(tm1978)* double mutant, the percentage of larvae with seam cell divisions was significantly different even after one day of L1 starvation (Fig. 4B).

**Fig. 4.**
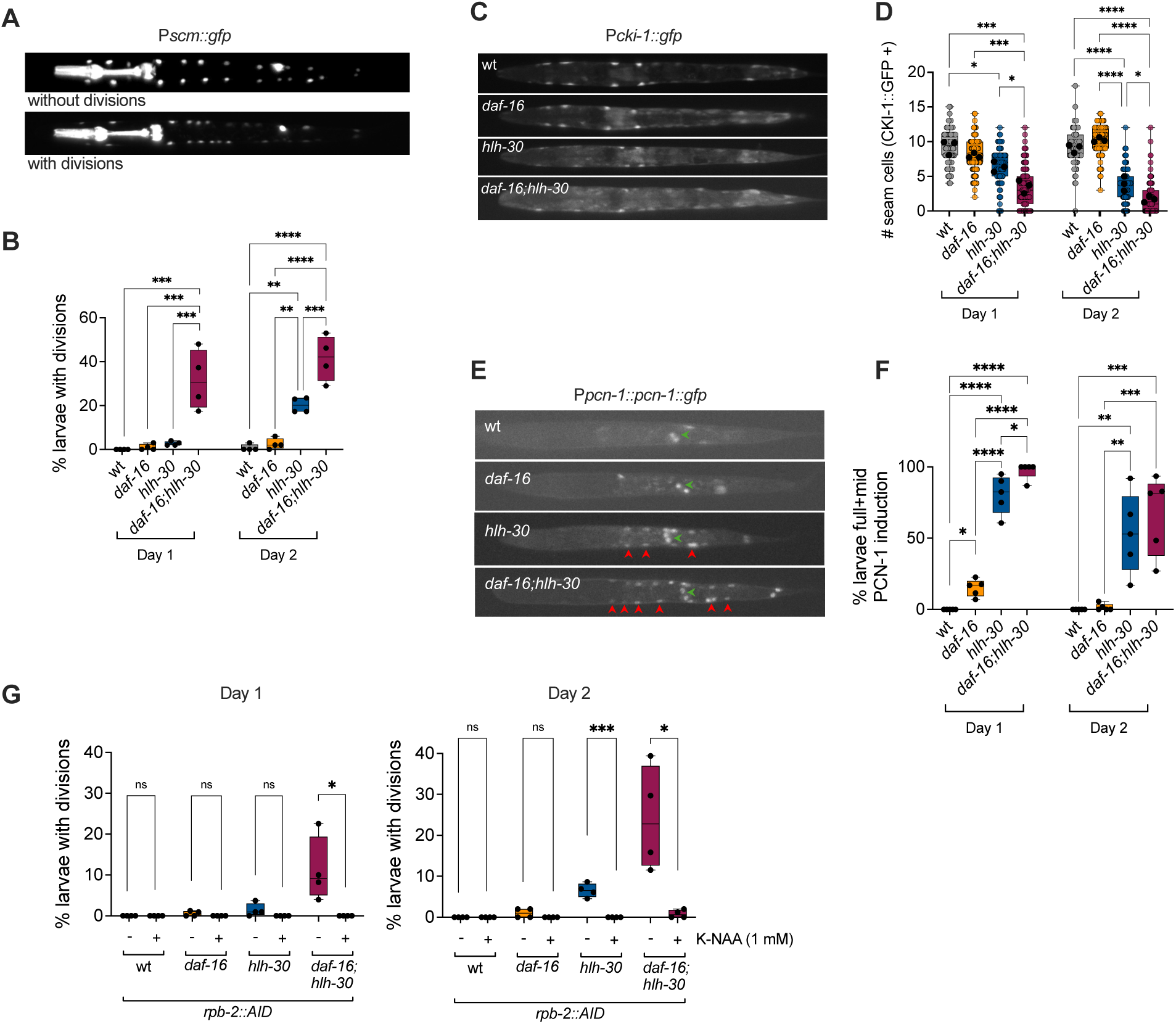
HLH-30 is necessary for cell cycle arrest during starvation. **(A)** Representative images of larvae without or with seam cell divisions. **(B)** Percentage of larvae with seam cell divisions after 1 or 2 days of L1 starvation, for the wild type, *daf-16(mu86)*, *hlh-30(tm1978)*, and *daf- 16;hlh-30*. Each dot represents the values for one independent experiment. **(C)** Representative images of *cki-1* expression induction during L1 starvation, for the wild type, *daf-16(mu86)*, *hlh- 30(tm1978)*, and *daf-16;hlh-30*. **(D)** Number of seam cell per larvae with induced *cki-1* (CKI*-* 1::GFP+) after 1 or 2 days of L1 starvation, for the wild type, *daf-16(mu86)*, *hlh-30(tm1978)*, and *daf-16;hlh-30*. Colored circles show values for individual larvae and black dots show the average for each experiment. **(E)** Representative images of *pcn-1* expression during L1 starvation, for the wild type, *daf-16(mu86)*, *hlh-30(tm1978)*, and *daf-16;hlh-30*. Red arrowheads point to expression in seam cell and green arrowheads to expression in Z2/Z3. **(F)** Fraction of larvae with different mid or high levels of PCN-1 induction after 1 or 2 days of L1 starvation, for the wild type, *daf-16(mu86)*, *hlh-30(tm1978)*, and *daf-16;hlh-30*. **(G)** Percentage of larvae with seam cell divisions after 1 or 2 days of L1 starvation, for the wild type, *daf-16(mu86)*, *hlh-30(tm1978)*, and *daf-16;hlh-30*, in control conditions (-) or treated with for the Auxin analogue NAA (+) for degradation of the RNA Polymerase II subunit *rpb-2*. Each dot represents the values for one independent experiment. In B, D and F, we performed One-way ANOVA followed by Tukeýs multiple comparisons. Only significant differences are shown. In G, we used *t-test* to evaluate the effect of *rpb-2* degradation on each genetic background. All plots are box and whiskers representing the independent replicates, marked with black dots, except for D, where the box and whiskers show the values for individual larvae and the averages per experiment are marked by black dots.

Previous research demonstrated that DAF-16 is necessary for cell cycle arrest during L1 starvation ^26^, but only in the presence of ethanol in the arrest media ^25^. These findings suggested that ethanol may suppress other factors that function redundantly with DAF-16 to maintain cell cycle arrest under starvation conditions. Given our evidence supporting a critical role for HLH-30 in this process, we investigated whether ethanol affects its subcellular localization. Notably, ethanol addition significantly reduced the nuclear localization of HLH-30 during L1 starvation (Fig. S5A). This suggests that ethanol-mediated deactivation of HLH-30 may enhance the relative importance of DAF-16 in sustaining cell cycle arrest during nutrient deprivation, explaining why the role of HLH-30 in cell cycle arrest had passed unnoticed.

To confirm the impact of HLH-30 on cell cycle regulators, we checked the expression of the cyclin-dependent kinase inhibitor CKI-1/CIP/KIP/p27, which increases over time during L1 starvation ^11,12^. We tested the induction of *cki-1* in the wild type and mutants by quantifying the number of seam cells with induced *Pcki-1::gfp* per larvae (Fig. 4C). The mutants *hlh-30* and *daf-16;hlh-30* showed a significant reduction in the number of seam cells (per larvae) with induced *Pcki-1::gfp*, compared to the wild type and the *daf-16* mutant (Fig. 4D). Finally, we tested the abundance of the S-phase marker PCN-1 using a translational reporter of PCN-1 (Fig. 4E). In all strains, the signal from the reporter was evident on Z2/Z3 PGCs, as these cells arrest at the G2 phase of the cycle during L1 starvation. The GFP signal was absent from other cells in the wild-type strain and showed non-significant levels of activation in the *daf-16* mutant on day 1. The *hlh-30* and *daf-16(mu86); hlh-30(tm1978)* mutants, however, showed significantly higher levels of activation of this marker in seam cells (Fig. 4F, Fig. S5B). *pcn-1* RNA is also more abundant in *hlh-30* and *daf-16;hlh-30* mutants (Fig. S5C). All these results are consistent with defective inhibition of the cell cycle in *hlh-30* mutants.

To further prove that deregulation of mRNA transcription underlies aberrant cell cycle activity in *hlh-30* mutants, we tested whether RNA polymerase II activity is required for these inappropriate divisions. We tested the effect of degradation of the polymerase II subunit RPB- 2 on cell division. For specific RPB-2 degradation, we used a strain with a tag for auxin induced degradation (AID) at the endogenous *rpb-2* locus ^27^ and checked the fraction of L1 with divisions after 1 and 2 days of starvation. The aberrant seam cell divisions in the *hlh-30* and *daf-16;hlh-30* mutants during starvation could be rescued by degradation of the RNA polymerase II subunit *rpb-2* (Fig. 4G). Together, these findings reveal a previously unappreciated role for HLH-30 in maintaining cell cycle quiescence during L1 starvation, acting in parallel to DAF-16 and potentially through modulation of RNA polymerase II activity.

### DAF-16 and HLH-30 reciprocally impact their function in regulation of gene expression

To find candidates for the effect of HLH-30 on cell cycle progression we turned again to the mRNA-seq results. The elevated number of differentially regulated genes in the *hlh-30* mutant makes it complex to tackle this question. Considering that changes in the activity of a single TF can provoke the differential expression of hundreds of genes, we rather analyzed transcription factor (TF) overactivation in the mutants. For this task, we used CelEst, an unified gene regulatory network built to estimate the activity of 487 distinct *C. elegans* TFs based on differential genes expression ^28^. As a result, we obtained a score for the activity of each TF in each of the mutants, relative to the wild type. In the *daf-16* mutant, the top overactivated TF is HLH-30 (Fig. 5A; File S1), suggesting that in the absence of DAF-16, HLH-30 plays a more prominent role in L1 starvation. This result suggests that the increased HLH-30 nuclear localization in the *daf-16* mutant (Fig. 1C) confers a functional difference, as it impacts expression of HLH-30 target genes. Other TFs overactivated in the *daf-16* mutant are CEY-1, SMA-9, MADF-10 and TLP-1 (Fig. 5A; File S1). Focusing on the TFs with reduced activity relative to the wild type, besides the obvious DAF-16 itself, we find that EFL-1, PHA-4, FKH-9 and HIF- 1, among others.

**Figure 5.**
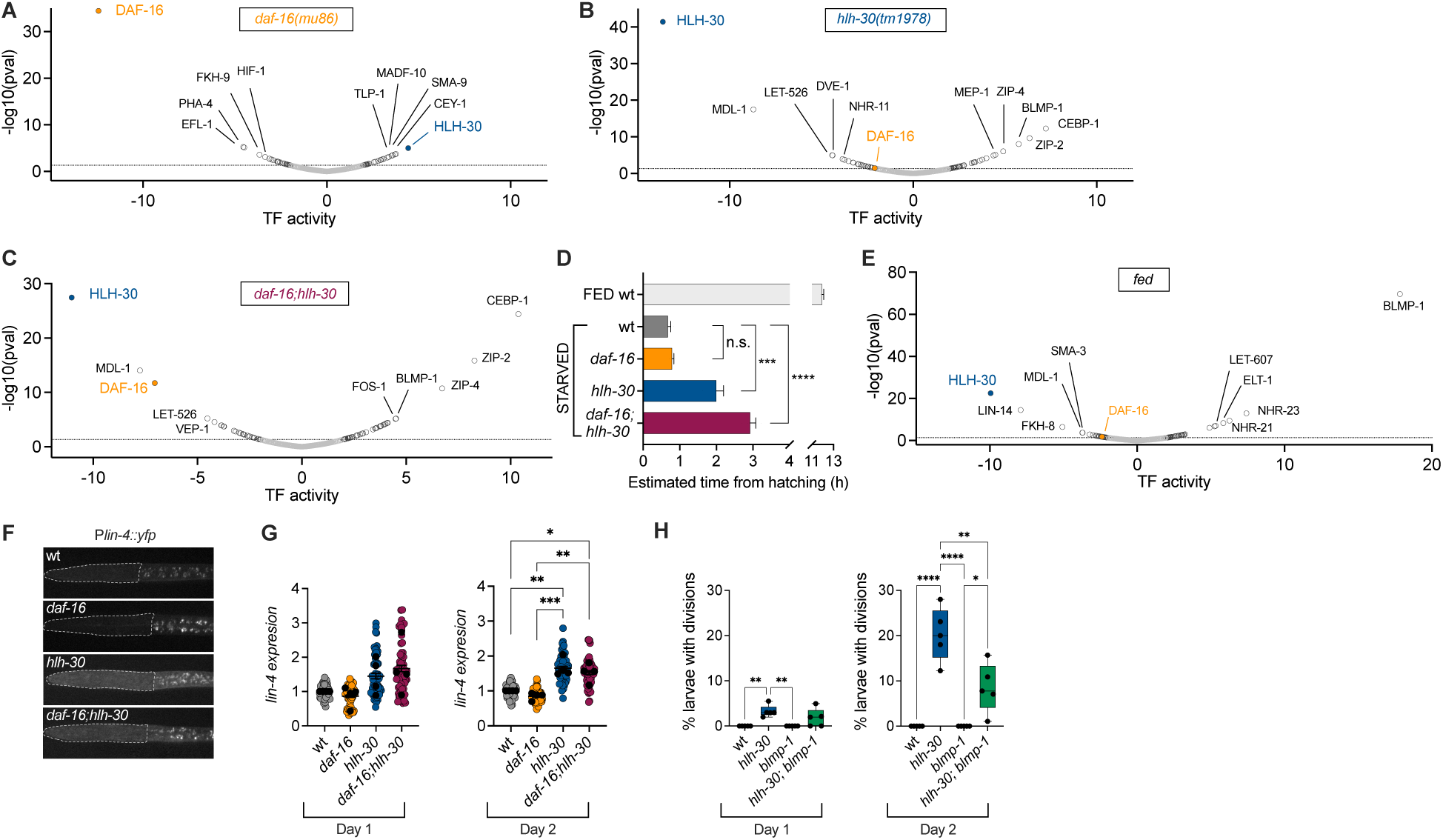
*hlh-30* mutation leads to BLMP-1 activation. (A-C) Plots showing TF activity score relative to the wild type for day-1 starved larvae of the *daf-16* mutant (A), *hlh-30* (B) and *daf- 16;hlh-30* (C). In A-C, the dotted lines at y=1.333 marks the significance threshold. TF activity changes over the threshold are represented by black circles and below are represented by grey circles. **(D)** Estimated age from hatching calculated by RAPToR, based on the overall gene expression for each condition. Error bars represent SEM. We compared the averages of each strain using One-way ANOVA followed by Dunnett’s test to compare each sample to the wild- type strain. **(E)** Plot showing TF activity score for fed versus starved wild type larvae. **(F)** Representative pictures showing *lin-4* expression during L1 starvation, for the wild type, *daf- 16(mu86)*, *hlh-30(tm1978)*, and *daf-16;hlh-30*. **(G)** Normalized expression levels of *lin-4::yfp* after 1 or 2 days of L1 starvation, for the wild type, *daf-16(mu86)*, *hlh-30(tm1978)*, and *daf- 16;hlh-30*. Colored circles show values for individual larvae and black dots show the average for each experiment. Lines mark the mean (±SEM). **(H)** Percentage of larvae with seam cell divisions after 1 or 2 days of L1 starvation, for the wild type, *hlh-30(tm1978)*, *blmp-1(s71)* and *hlh-30;blmp-1*. Plots are box and whiskers representing the independent replicates, marked with black dots. In G and H, for statistical analysis, we performed One-way ANOVA followed by Tukeýs multiple comparisons. Only significant differences are shown.

Mutation of *hlh-30* reduced the activation of DAF-16 (p-value 0.036) relative to the wild type (Fig. 5B; File S1), a result that also agrees with our observation at the level of subcellular localization (Fig. 1B). In the *hlh-30* mutant the top overactivated TFs are CEBP-1, ZIP-2, BLMP- 1, ZIP-4 and MEP-1. The TFs with reduced activity in this mutant are HLH-30, MDL-1, LET-526, DVE-1 and NHR-11. In the double mutant, the lists over activated and down activated TFs largely overlapped with those of the *hlh-30* mutant (Fig. 5C; File S1).

We also investigated the effect of ethanol in TF activation in the *daf-16* mutant. We used an available dataset of differentially expressed genes between the *daf-16* mutant and the wild type under L1 starvation in S-basal with 0.1% ethanol ^29^. Under these conditions, the *daf-16* mutant does not exhibit the HLH-30 overactivation that we observed in complete starvation (Fig. S6A). Additionally, we calculated the correlation between TF activity scores in the *daf-16* mutant with ethanol and those in the single and double mutants without ethanol, across all possible combinations. We found that the *daf-16* mutant without ethanol showed a mild negative correlation with *hlh-30*, likely due to the overactivation of HLH-30 targets in the *daf-16* mutant, and no correlation with the *daf-16;hlh-30* double mutant (Fig. S6A,C). In contrast, the *daf-16* mutant exposed to ethanol displayed a significant positive correlation with both *hlh-30* and *daf-16;hlh-30* (Fig. S6B). This result suggests that the reduced nuclear localization of HLH-30 in the presence of ethanol (Fig. S5A) has functional consequences, specifically dampening the transcriptional activation of HLH-30 target genes. In other words, ethanol masks the role of HLH-30 during L1 arrest by deactivating this transcription factor. Instead, in complete starvation these two transcription factors collaborate and reciprocally impact their activation.

### BLMP-1 is overactivated in *hlh-30* mutants and contributes to initiate development under starvation

Considering the impaired seam cell arrest of the *hlh-30* and *hlh-30;daf-16* mutants, we decided to compare TF activation in these mutants with that of developing L1 larvae. We performed another mRNA-seq experiment to assess gene expression and TF activation in developing (fed) L1 compared to arrested (starved) L1. When we compared TF activation in the mutant samples among each other, and to that of the fed sample, we observed that *hlh-30* and *daf-16;hlh-30* mutants showed a significant positive correlation with the fed condition (Fig. S6C). Additionally, *hlh-30* and *daf-16;hlh-30* showed the strongest correlation between each other. However, the relative TF activation of the *daf-16* mutant did not correlate with that of the fed sample and showed a weak negative correlation with the *hlh-30* mutant (Fig. S6B,C). This result suggest that the *hlh-30* mutation confers transcriptional features that are similar to that of fed developing larvae.

With these clues, we decided to use RAPToR (Real Age Prediction from Transcriptome staging On Reference). RAPToR uses interpolation of gene expression time-series to predict the developmental age of a sample from their expression profile ^33^. With this analysis, our fed L1 sample showed an estimated age of approx. 12 hours, consistent with samples been collected 24 hours after embryo preparation (Fig. 5D). The estimated time from hatching of the starved wild-type and *daf-16* mutant samples was less than 1 hour, consistent with developmental arrest after hatching. However, *hlh-30* and *daf-16;hlh-30* showed an advanced developmental stage significantly different to that of the wild type (Fig. 5D).

Interestingly, we observed that HLH-30 was the most inactivated TF and BLMP-1 the highest overactivated TF in fed larvae (Fig. 5E, File S1). BLMP-1 expression and priming activities are normally repressed during L1 arrest and expressed in animals that have been re- fed. This way, BLMP-1 acts as a pioneer TF to decompact the *lin-4* locus in response to nutrients before transcriptional activation throughout development ^30^. The initiation of larval development involves the activation of the transcription of the miRNA *lin-4* ^31^. In the wild-type strain, *lin-4* transcription is not observed in starvation-arrested L1 larva ^32^. When we check *blmp-1* expression from the mRNA-seq data, we observed that, indeed, *blmp-1* expression was significantly increased in the *hlh-30* mutant and the double mutant, compared to the wild type and *daf-16* (Fig. S6D). Indeed, BLMP-1 appears among the list of top 5 overactivated TFs in the *hlh-30 and the hlh-30;daf-16* mutants (Fig. 5B-C). Then, we investigated the expression of the microRNA *lin-4* using a *Plin-4::yfp* reporter strain (Fig. 5F). We observed that the levels of the miRNA *lin-4* were increased in the *hlh-30* and double mutants, being significantly different from the wild type and the *daf-16* mutant after two days of L1 starvation (Fig. 5G).

All this information makes BLMP-1 an interesting candidate to mediate arrest-defective phenotype of *hlh-30*, as the overactivation of BLMP-1 could activate the developmental program during starvation in this mutant. To test this hypothesis, we analyzed seam cell divisions in wild-type, *hlh-30,* the *blmp-1(s71)* and *hlh-30;blmp-1* backgrounds. We observed that the *blmp-1* mutation partially rescues the increase in seam cell divisions caused by HLH-30 loss (Fig. 5H). These results support a model in which HLH-30 ensures proper developmental arrest by repressing BLMP-1 activity under starvation conditions, thereby preventing premature transcriptional priming and cell cycle progression.

## DISCUSSION

### Functional collaboration between DAF-16 and HLH-30

The transcription factors DAF-16/FOXO and HLH-30/TFEB frequently act together in *C. elegans*, forming a combinatorial transcriptional module that responds to various stressors, including oxidative stress, starvation, and aging. This synergy is stimulus-dependent and reflects a finely tuned evolutionary adaptation to environmental challenges ^18,34^. Both DAF-16 and HLH-30 are critical regulators that support L1 survival during starvation. DAF-16 and HLH- 30 share a substantial number of transcriptional targets—approximately 64% (1,894 out of 2,975) of the genes differentially regulated in *daf-16* mutants are also misregulated in *hlh-30* mutants (Fig. 2B).

In this study, we examined their functional interdependence during L1 arrest and observed a reciprocal influence in their activation. Notably, in the absence of DAF-16, HLH-30 becomes overactivated during L1 arrest. This conclusion is supported by both subcellular localization data and the upregulation of HLH-30 target genes in *daf-16* mutants (Fig. 1B and Fig. 5A). Conversely, *hlh-30* mutants do not exhibit overactivation of DAF-16. Instead, they show reduced activation of this transcription factor (Fig. 1A and Fig. 5B). Interestingly, the hypomorphic *daf-2(e1370)* mutation enhances survival and recovery in *hlh-30* mutants in a DAF-16-dependent manner, implying that nuclear DAF-16 may partially substitute for HLH-30 function. DAF-16 and HLH-30 are known to physically interact under stress conditions in other contexts ^18^, and both localize to the nucleus at day 1 of L1 arrest. However, the dynamic patterns of their nuclear localization during L1 arrest (Fig. 1A–B) suggest that their interaction in this specific context warrants further investigation.

### HLH-30 as a master regulator during L1 starvation

Previous investigations into the role of HLH-30 during L1 arrest have primarily focused on its function as a regulator of lysosome biogenesis. In particular, HLH-30–dependent activation of the lysosomal lipase *lipl-2* has been shown to be essential for survival under starvation conditions ^16^. In this study, we reveal additional, previously unrecognized roles for HLH-30 during L1 starvation. First, HLH-30 is required for chromatin reorganization in response to nutrient deprivation (Fig. 3A–C). Under starvation, chromatin in the intestine undergoes a distinct reorganization, forming two concentric rings at the nuclear and nucleolar periphery. This structural change is mediated by the inactivation of mTOR signaling ^13^ and we now demonstrate that HLH-30, as an mTOR signaling effector, is necessary for this reorganization.

Most notably, HLH-30 is also required for cell cycle arrest during L1 starvation (Fig. 4), a finding that was unexpected given that this role had previously been attributed primarily to DAF-16 ^26^. We have unveiled that this discrepancy likely stems from differences in experimental conditions. In earlier studies demonstrating DAF-16’s role in activating *cki-1* and promoting cell cycle arrest, the starvation medium contained ethanol. Subsequent research revealed that the failure of *daf-16* mutants to arrest cell divisions was ethanol-dependent and did not occur in ethanol-free M9 buffer ^25^. This suggests that ethanol interferes with alternative compensatory mechanisms that might otherwise suppress proliferation in the absence of DAF-16. Our data further support this interpretation: we observed that ethanol reduces HLH-30 nuclear localization (Fig. S5A) and target gene activation (Fig. S6A), indicating that ethanol may directly impair HLH-30 function. L1 larvae can use ethanol as a carbon source promoting the synthesis of fatty acids and amino acids, that potentially can activate the kinase activity of TORC1 ^35,36^. These findings suggest that DAF-16 and HLH-30 act redundantly to mediate cell cycle arrest during L1 starvation. However, in the presence of ethanol, HLH-30 is deactivated, rendering DAF-16 essential for this process. Consistent with this, *hlh-30* mutants exhibit defective cell cycle arrest, likely due to the combined absence of HLH-30 and the reduced activation of DAF-16 in these mutants.

### *hlh-30* mutants show transcriptional features of developing larvae

Our data reveal that *hlh-30* mutants fail to arrest cell divisions during L1 starvation and instead exhibit a transcriptional profile resembling that of fed, developing larvae (Fig. 5G, S6C). A defining feature of this developmental transcriptional state is the activation of BLMP-1 target genes in fed larvae (Fig. 5D), which are also markedly overexpressed in starved *hlh-30* and *daf-16;hlh-30* mutants (Fig. 5B-C). This suggests that in the absence of HLH-30, the starvation-induced developmental arrest program is compromised, allowing inappropriate activation of developmental pathways. Notably, mutation of *blmp-1* partially rescues the defective cell cycle arrest phenotype of *hlh-30* mutants (Fig. 5H), indicating that BLMP-1 overactivation contributes directly to the failure to suppress proliferation. These findings support a model in which HLH-30 represses BLMP-1-driven transcriptional programs during starvation, ensuring that larvae enter and maintain a quiescent state.

BLMP-1 is a conserved transcription factor that coordinates post-embryonic developmental programs and modulates the transcriptional output of cyclically expressed genes, contributing to developmental robustness in *C. elegans* ^37,38^. It is homologous to mammalian BLIMP-1 (PRDM1), a key regulator of cell fate decisions (reviewed in ^39^). Recent work has shown that BLMP-1 facilitates transcriptional priming by promoting chromatin decompaction at target loci, thereby modulating the amplitude and duration of gene expression without altering its timing. This chromatin remodeling is regulated by nutritional inputs and becomes essential when animals resume development after nutrient-mediated arrest ^30^. In *hlh-30* mutants, the overactivation of BLMP-1 targets during starvation likely reflects a failure to suppress this priming mechanism, leading to inappropriate cell cycle progression. Together, our results highlight the critical role of HLH-30 in integrating environmental cues with developmental timing by antagonizing BLMP-1 activity, thereby safeguarding the transcriptional integrity of the starvation-induced arrest program.

### Broader Implications for cancer and cell senescence

Transcription Factor EB (TFEB) is a central regulator of cellular homeostasis in mammals, primarily recognized for its role in controlling autophagy and lysosomal biogenesis. It activates genes involved in the degradation and recycling of cellular components, particularly under stress conditions such as nutrient deprivation or oxidative damage. Beyond these functions, TFEB contributes to metabolic regulation, neuroprotection, immune responses, and inflammation. Dysregulation of TFEB has been implicated in various diseases, including neurodegenerative disorders and several types of cancer (reviewed in ^40^). The rapid proliferation of cancer cells increases their demand for energy and biosynthetic materials, making enhanced autophagolysosomal activity a common feature in tumors and positioning TFEB as a promising target in cancer research. Interestingly, the oncogenic potential of TFEB appears to be more closely tied to its regulation of the cell cycle than to macroautophagy. For example, TFEB can transcriptionally upregulate CDK4/6, promoting the G1/S transition and cell division ^41^. Conversely, TFEB also exhibits tumor-suppressive properties. We have shown that HLH-30, its *C. elegans* ortholog, is required for CKI-1 induction during starvation, and in human cell lines such as HeLa and ARPE-19, TFEB depletion significantly reduces basal p21 expression. Overexpression of TFEB, on the other hand, upregulates p21 and induces G1 arrest ^42^. In respect to senescence, TFEB targets are highly expressed in quiescent cells and TFEB activation can boost lysosomal function of senescent cells to improve its reactivation in response to proliferative signals ^7^. These findings underscore the context-dependent nature of TFEB’s role in cancer, highlighting its dual capacity to either promote or suppress tumorigenesis. Our findings on the role of HLH-30/TFEB in cell cycle arrest underscore the role of *C. elegans* L1 starvation to understand cellular processes related to quiescence maintenance.

## MATERIALS AND METHODS

### *C. elegans* strains and growth conditions

Nematodes were grown according to standard methods, maintaining them on nematode growth medium (NGM) agar plates with *Escherichia coli* OP50-1 bacteria at 20 °C. All strains used in this study are listed in Supplementary Table 1.

### Generation of starved L1 larvae

To induce L1 larval arrest by starvation, embryos were isolated from gravid adults using an alkaline hypochlorite solution. The embryos were then resuspended in M9 minimum medium at a density of 20 embryos per microliter and were incubated at 20°C with gentle shaking, allowing them to hatch. In the absence of food, the newly hatched larvae arrested development at the L1 stage. We count the days of arrest from the moment of embryos preparation, that is, day 1 corresponds to 24 hours after embryo preparation.

### Generation of fed L1 larvae

For the fed L1 samples, we isolated embryos as detailed above but resuspended them in S-basal containing *E. coli* OP50-1 at 10 gl^-1^. In this condition, larvae initiate postembryonic development after hatching, that occurs approximately 12 hours after embryo preparation. Thus, after 24 hours, fed samples are approximately 12 hours into postembryonic development.

### Subcellular localization of HLH-30 and DAF-16

To determine the subcellular localization of HLH-30 and DAF-16, strains expressing the fluorescently tagged versions of these proteins (HLH-30::GFP and DAF-16::GFP) were used. At the indicated time points, 18 µl of the suspension of L1-arrested larvae were collected and incubated with 2 µl of 1 mM Levamisol (for HLH-30 localization) or 10 mM Levamisol (for DAF- 16). After precipitation, the number of animals exhibiting cytoplasmic, nuclear, or intermediate localization of these reporters was counted in a population of at least 100 animals per sample. Based on these data, the percentage of animals with each localization pattern was calculated. To prevent artificial translocation of HLH-30, a reduced concentration of Levamisole was used. Similarly, room temperature was maintained at 20°C during visualization to avoid potential temperature-induced effects on HLH-30 or DAF-16 activation. A minimum of three independent biological replicates for each time point were analyzed for each condition.

### Survival and recovery assays

To evaluate survival of starved L1 larvae, we pipetted 5 µl of day 1-arrested larvae onto a drop of M9 on a glass slide and scored them directly under a dissecting scope. We counted the number of live larvae and divided it by the total number of larvae observed to calculate the percentage of survival. To measure recovery capacity of the starved L1 larvae, we pipetted 5 μl of day 1-starved larvae to NGM plates with OP50-1. We counted the number of larvae and then incubated the plates at 20 °C. After 2 days, we counted the number of larvae that had resumed development and divided by the number of larvae counted after plating to calculate the percentage of larvae that recovered. At least three independent biological replicates were performed.

### Total RNA sample preparation and mRNA sequencing

We performed two complete experiments, each including biological replicates. Experiment 1 aims to evaluate the contribution of the transcription factors DAF-16 and HLH-30 to gene expression during L1 starvation. Samples consisted of L1 of the wild-type strain and the mutants *daf-16(mu86)*, *hlh-30(tm1978)* and *daf-16(mu86); hlh-30(tm1978)* collected on day 1 of L1 starvation in M9 buffer. We performed three independent biological replicates of this experiment. Experiment 2 aims to evaluate the impact of L1 starvation in gene expression. Samples of L1 larvae of the wild-type strain, either starved or fed, also collected at day 1 (24 hours from embryo isolation). We performed two independent biological replicates of this experiment. For RNA extraction, we collected at least 140,000 L1 larvae per sample. Total RNA was extracted using the NZYTotal RNA Isolation Kit (NZYtech, MB13402). A total of 150 ng of RNA per sample was used to prepare libraries following the Illumina Stranded mRNA Prep Ligation, compatible with the Illumina sequencing system. The quality of the libraries was assessed for adequate size distribution and integrity using the TapeStation DNA High Sensitive D1000 assay, and sufficient concentration using Qubit™ DNA HS Assay fluorimetry. Samples were sequenced using the Illumina NextSeq 500 High Output platform.

### RNA-seq functional analysis and visualization

Quality control and cleaning of the raw reads was performed with FASTP package (v0.22.0). Clean reads were then aligned to the *C. elegans* reference genome (https://ftp.ensembl.org/pub/release-110/gff3/caenorhabditis_elegans/) using the STAR package (v2.7.10b). Raw counts for individual genes were quantified with the featureCounts command of the SUBREAD(v.2.0.3) package. To normalize the counts, we applied TMM (trimmed mean of M values) normalization with edgeR (v3.34.1) package within the R environment. DEGs was analyzed in the mutant strains in comparison to the wild type. Pairwise comparisons were performed between each of the mutants and the wild-type strain using DESeq2 (v1.42.0) ^43^. The genes of the three conditions with an adjusted p-value <0.05 (Wald test), regardless of the Fold Change between samples, were selected for further analysis.

Functional enrichment analysis of DEGs was performed employing the clusterProfiler package (v4.14.6) ^22^, using Gene Ontology (GO). Ensembl gene IDs were converted to Entrez IDs prior to enrichment. GO enrichment analysis was performed using the enrichGO() function, specifying the biological process ontology (ont = “BP”) and applying a significance threshold of p.adjust <0.01, using the Benjamini-Hochberg method. Redundant terms were reduced using the simplify() function with a cutoff of 0.8. For visualization, we generated a scatter plot of the 20 most significant GO terms with ggplot2 (v3.5.2) ^44^. Gene expression patterns related to the mitotic cell cycle and chromosome organization were displayed as a heat map using ComplexHeatmap (v2.22.0) ^45^ (after Z-score scaling. GO gene networks were constructed with the cnetplot() function (enrichplot v1.26.6), retaining the 12 most significant terms per condition. Data preprocessing was performed with readxl (v1.4.5) and dplyr (v1.1.4).

### Analysis of chromatin spatial distribution

Microscopy was performed using a live-cell imaging system (Confocal Spinning Disk Microscope) from Visitron Systems GmbH, equipped as follows: Nikon Eclipse Ti2 microscope with a Plan Apo λ 100×/1.45 oil objective, Yokogawa CSU-W1 confocal scanner unit, VS-Homogenizer, Electron Multiplying CCD camera (Andor iXon Series), and VisiView software for image acquisition. For the analysis of chromatin profiling in intestinal nuclei, the protocol previously described ^13^, was followed. In brief, live imaging was conducted on 2% agarose pads supplemented with 0.15 % sodium azide to immobilize the worms. All images were acquired with 50 stacks and a z-spacing of 200 nm. The nuclei of the mid-intestine were manually segmented in 2D, based on HIS-72::GFP, and the nucleolar center annotated using Cell-ACDC^46^. To extract the intensity profiles, we proceed as described in ^13^. A total of 78 cells per strain were analyzed from three independent biological replicates.

### Fluorescence reporter analysis

For analyzing fluorescence reporters, unless otherwise specified, at the indicated time points, 50 µl of the suspension of L1-arrested larvae transferred to a clean tube and centrifuged for one minute at 850 g. Then, 40 µl of the supernatant were removed, and Levamisole was added to the sample at 10 mM to paralyze the animals. The sample was then transferred to a glass slide, covered with a coverslip, and the GFP signal was visualized using a Leica scope M205 FCA with GFP excitation/emission filters. Lin-4::YFP signal from each larva was normalized to the average signal from the wild-type strain in each replicate.

### Analysis of seam cell divisions

For analyzing seam cell divisions, we used strains carrying the *Pscm::NLS-GFP* construct, which is specifically expressed in seam cells. We counted the fraction of animals presenting one or more divided cells in the V1–V4 seam cell lineage. At least 50 larvae per strain and replicate were analyzed for each sample. We performed four independent biological replicates.

### Analysis of *cki-1* activation

For the analysis of *cki-1* activation, strains carrying the transcriptional reporter *cki-1::GFP* were used. Images of at least 80 larvae were acquired using fixed intensity and exposure parameters. Using FiJi/ImageJ, images were binarized. After binarization, the number of seam cells displaying signal per larva was quantified. Three independent biological replicates were performed. The GFP signal was visualized using a Leica scope M205 FCA with GFP excitation/emission filters.

### Analysis of PCN-1 induction

For the analysis of *pcn-1* activation, strains carrying the translational reporter *Ppcn-1::pcn- 1::GFP* were used. We counted the fraction of animals in a population of at least 50 larvae showing each category of PCN-1 induction (full, mid or no induction) as follows. Full induction corresponds to animals showing GFP signal in H and V1-V4 seam cells, mid induction corresponds to larvae showing GFP signal in at least on V1-V4 seam cell, and the fraction of animals not showing GFP signal was categorized as larvae without PCN-1 induction. We performed five independent biological replicates. The GFP signal was visualized using a Leica scope M205 FCA with GFP excitation/emission filters.

### Analysis of transcription factor activation and age estimates

To estimate transcription factor activation we used CelEsT (github.com/IBMB-MFP/CelEsT- app) ^28^. We used the raw count (File S1) from the mRNA-seq experiments to calculate the TF activation scores and p-values and represented them in Volcano plots. Within the app, also starting from raw counts, we used the RAPToR (Real Age Prediction from Transcriptome staging On Reference) package to estimate the age of each sample according to their expression data ^33^.

### Statistical analysis

For statistics, we have used the averages of independent biological replicates to avoid the inflated N value from using individual animals ^47^. A complete report of biological replicates and, when relevant, number of larvae per sample, can be found at Supplementary Table 2. We used unpaired two-tailed t-test to compare the means of two groups. For the rest of the results, we used one-way ANOVA to compare more than two groups. After one-way ANOVA, we either perform Dunnett’s multiple comparison to compare the average of each genotype to the average of the wild type, or Tukeýs multiple comparison to compare the average of each genotype to all the other genotypes. The type of analysis that applied to each panel is detailed in the figure legends. For clarity, some plots show only relevant comparisons. In all cases, * means p>0.05, ** p>0.01, *** p>0.001, and **** p>0.0001. Graphs and statistics were performed on Graphpad Prism 10.

### Data availability

All raw sequencing data for mRNA-seq libraries will be deposited at NCBI Gene Expression Omnibus (GEO). The mRNA-seq processed data (raw counts, normalized TMM, DESeq lists, TF activation scores) are included as Supplementary material (File S1).

## Acknowledgments

We thank Antonio Miranda-Vizuete (IBIS) and David Aristizabal (IDIBELL) for helpful discussion. Some strains were provided by the *Caenorhabditis* Genetics Center (CGC), which is funded by NIH Office of Research Infrastructure Programs (P40 OD010440). This work is supported by the grants PID2022-139009OB-I00 and PID2019-104632GB-I00 funded by MICIU/AEI/ 10.13039/501100011033, awarded to MO. MMB was supported by a postdoctoral contract from the “Programa de Ayudas a la I+D+i, en Régimen de Concurrencia Competitiva en el Ámbito del Plan Andaluz de Investigación, Desarrollo e Innovación (PAIDI 2020)”, and her work at the laboratory of DSC was supported by an EMBO “Scientific Exchange Grants”. AMC was supported by a contract of the VII PPIT- University of Sevilla (Contratos de Acceso al Sistema Español de Ciencia, Tecnología e Innovación). DSC thanks Helmholtz Munich for Support and the German Research Foundation (Deutsche Forschungsgemeinschaft) for the 2202 Priority Programme ‘Spatial Genome Architecture in Development and Disease’ CA 2753/2-1 and the individual grants CA2753/1-1 and CA2753/3-1.

## Supplementary information

**Figure S1.**
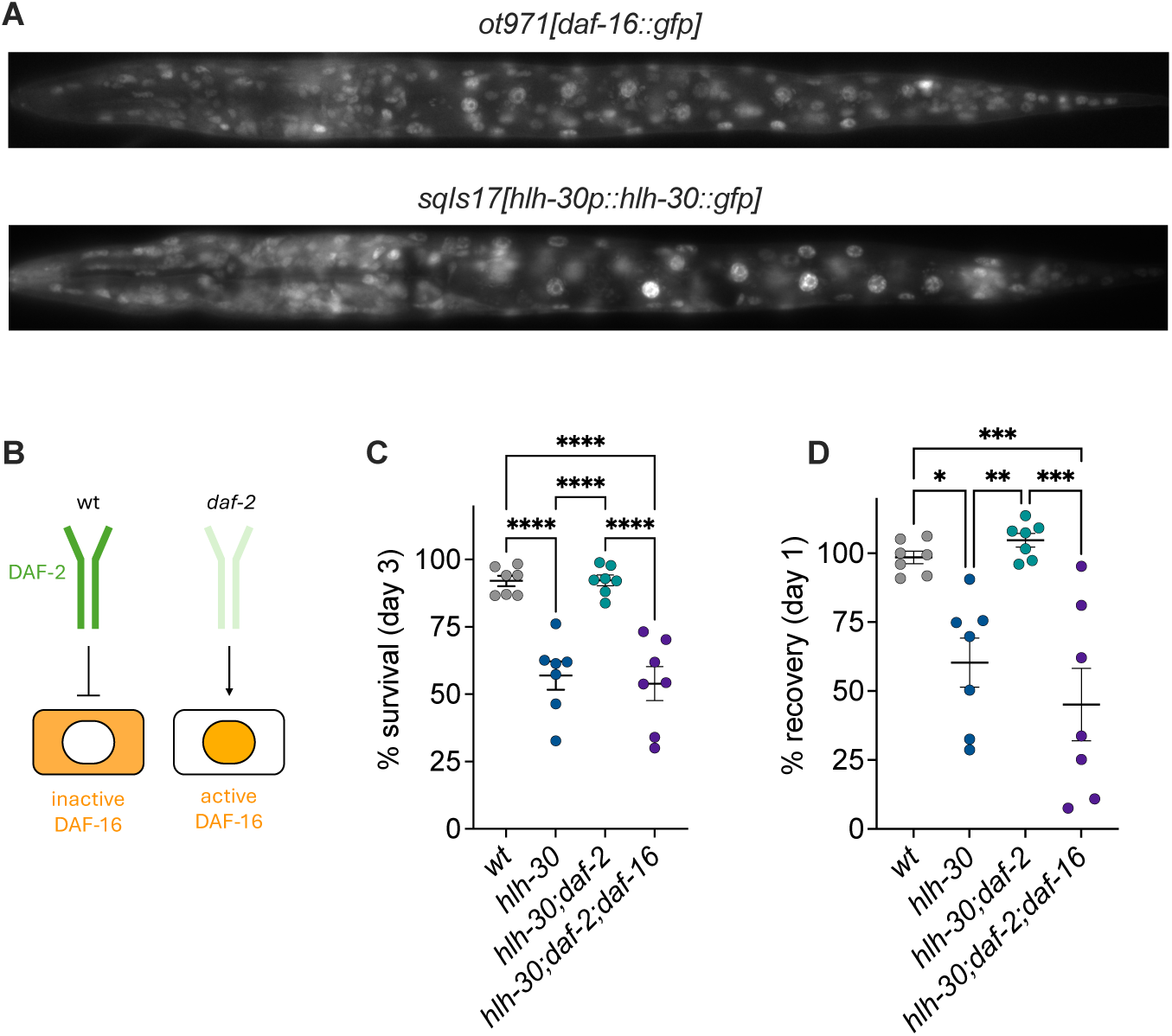
Relative to Figure 2. **(A)** Representative images of *ot971[daf-16::gfp]* and *sqIs17[hlh-30p::hlh-30::gfp]* larvae with strong nuclear localization. **(B)** Control of DAF-16 subcellular localization by insulin signaling. **(C)** Percentage of L1 larvae alive after 3 days of starvation for the wild-type, *hlh-30(tm1978)*, *hlh-30(tm1978);daf-2(e1370)* and *hlh-30(tm1978);daf-2(e1370);daf-16(mu86)*. **(D)** Percentage of L1 larvae that resume development upon refeeding after 1 day of starvation, for the same strains as in B. Colored dots represent results from independent experiments, and the lines mark the mean (±SEM). In (C) and (D), we performed One-way ANOVA followed by Tukeýs multiple comparisons to compare all samples. Only significative differences are shown. Complete statistics report can be found in File S1.

**Figure S2.**
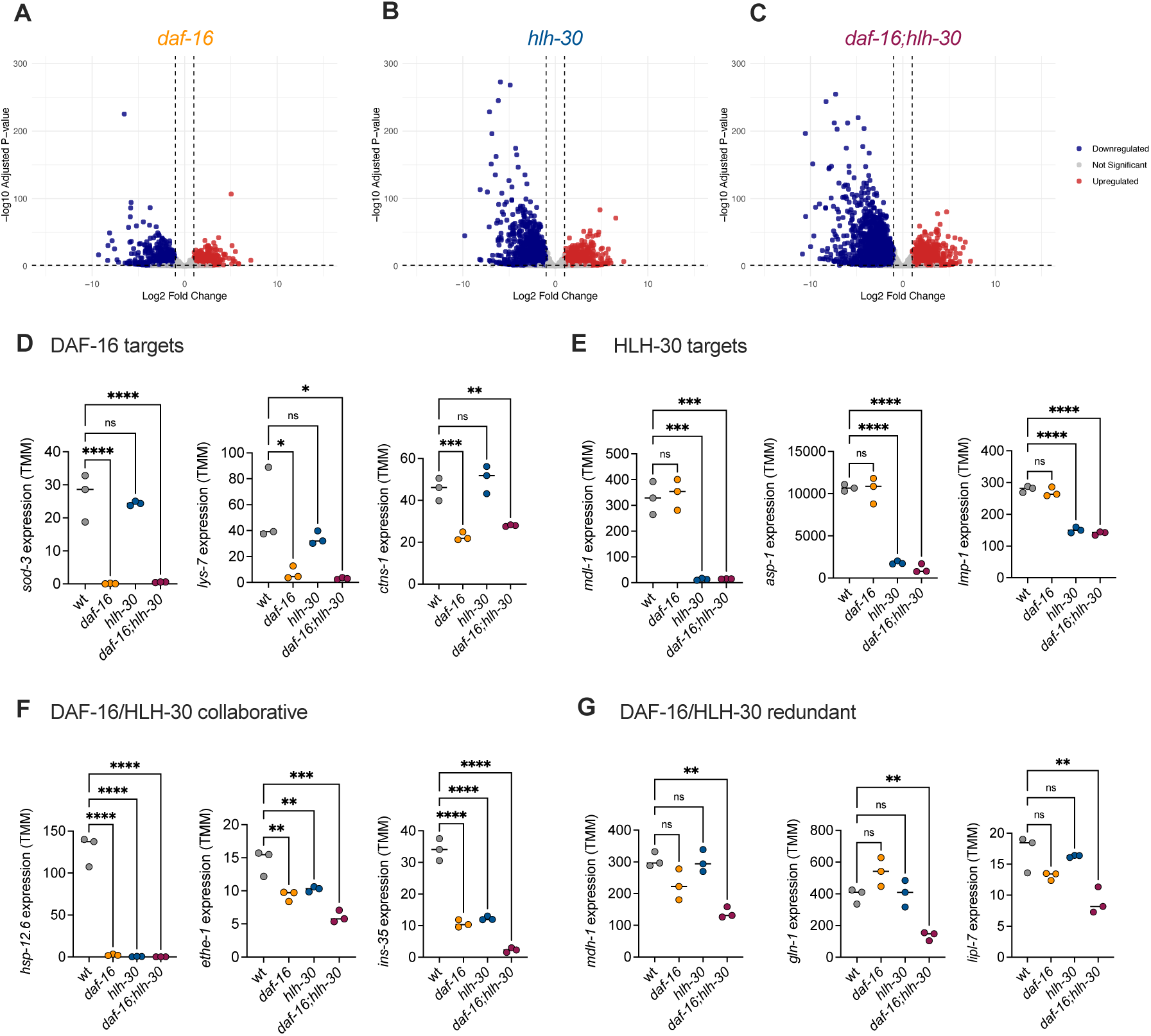
Relative to Figure 2. **(A-C)** Volcano plots showing Fold change and p-vale for all gene expression relative to the wild type. **(D)** Expression of the DAF-16 targets *sod-3*, *lys-7* and *ctns-1* in the wild-type, and the mutants *daf-16*, *hlh-30*, and the double mutant *daf-16*; *hlh-30*. **(E)** Expression of the HLH- 30 targets *mdl-1*, *asp-1* and *lmp-1* in the wild-type, and the *daf-16*, *hlh-30*, and *daf-16*; *hlh-30* mutants. **(F)** Expression of the genes *hsp-12.6*, *ethe-1* and *ins-35* in the wild-type, and the *daf-16*, *hlh-30*, and *daf-16*; *hlh-30* mutants. **(G)** Expression of the genes *mdh-1*, *gln-1* and *lipl- 7* in the wild-type, and the *daf-16*, *hlh-30*, and *daf-16*; *hlh-30* mutants. For D-G, results are shown as TMM values (trimmed mean of M; File S2) from the mRNA-seq experiment. Colored dots represent results from independent experiments, and the lines mark the mean. We performed One-way ANOVA followed by Tukeýs multiple comparison to assess the significance of the differences between each pair of strains. For clarity, the plots show only the comparisons of each mutant to the wild type. Complete statistics report can be found in File S1.

**Figure S3.**
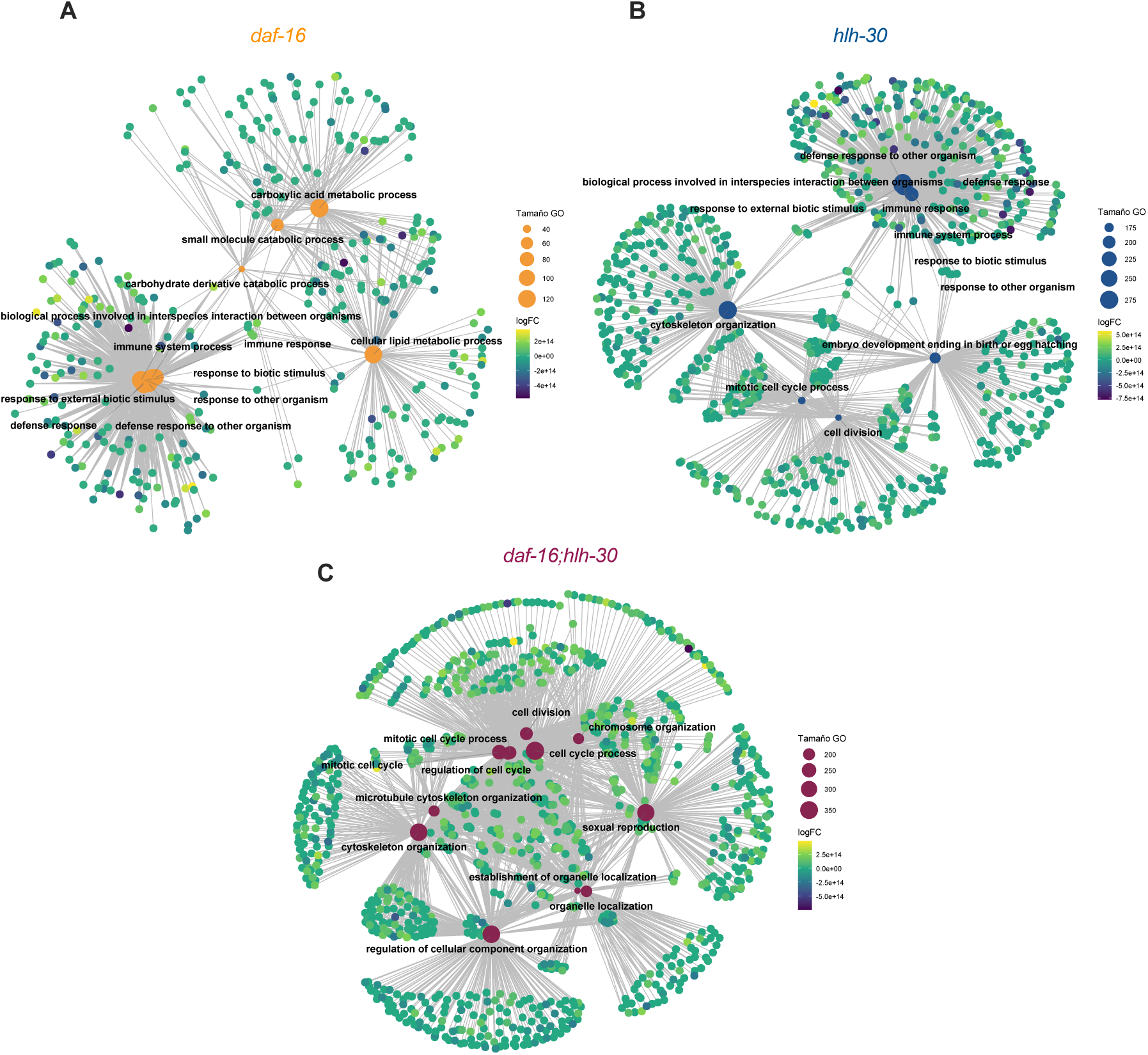
Relative to Figure 2. GO gene networks. The 12 most significant GO terms were used to generate the networks for *daf-16* **(A)**, *hlh-30* **(B)** and *daf-16; hlh-30* **(C)** mutant strains.

**Figure S4.**
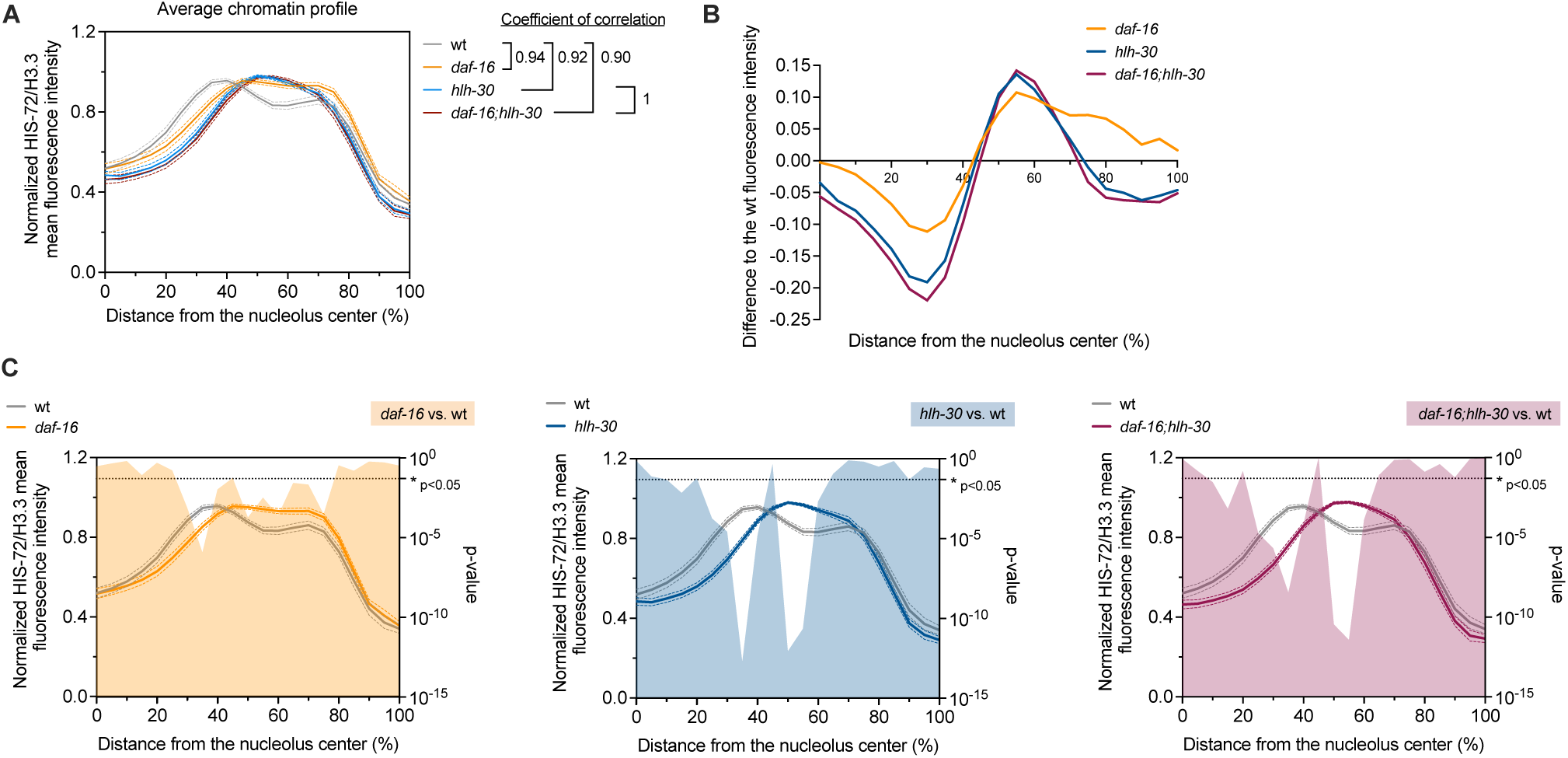
Relative to Figure 3. **(A)** Overlapped average chromatic prolife of the four strains represented in Fig. 3C. **(B)** Difference in fluorescence intensity between each of the mutants and the wild type, as a function of the distance from the nucleolus center. **(C)** Comparison of chromatin profiles for each mutant (as in Fig. 4C) overlapped with the results from t-test (right Y-axis) comparing the two profiles position by position from the nucleolus center (0 %) to the nuclear periphery (100 %). Complete statistics report can be found in File S1.

**Figure S5.**
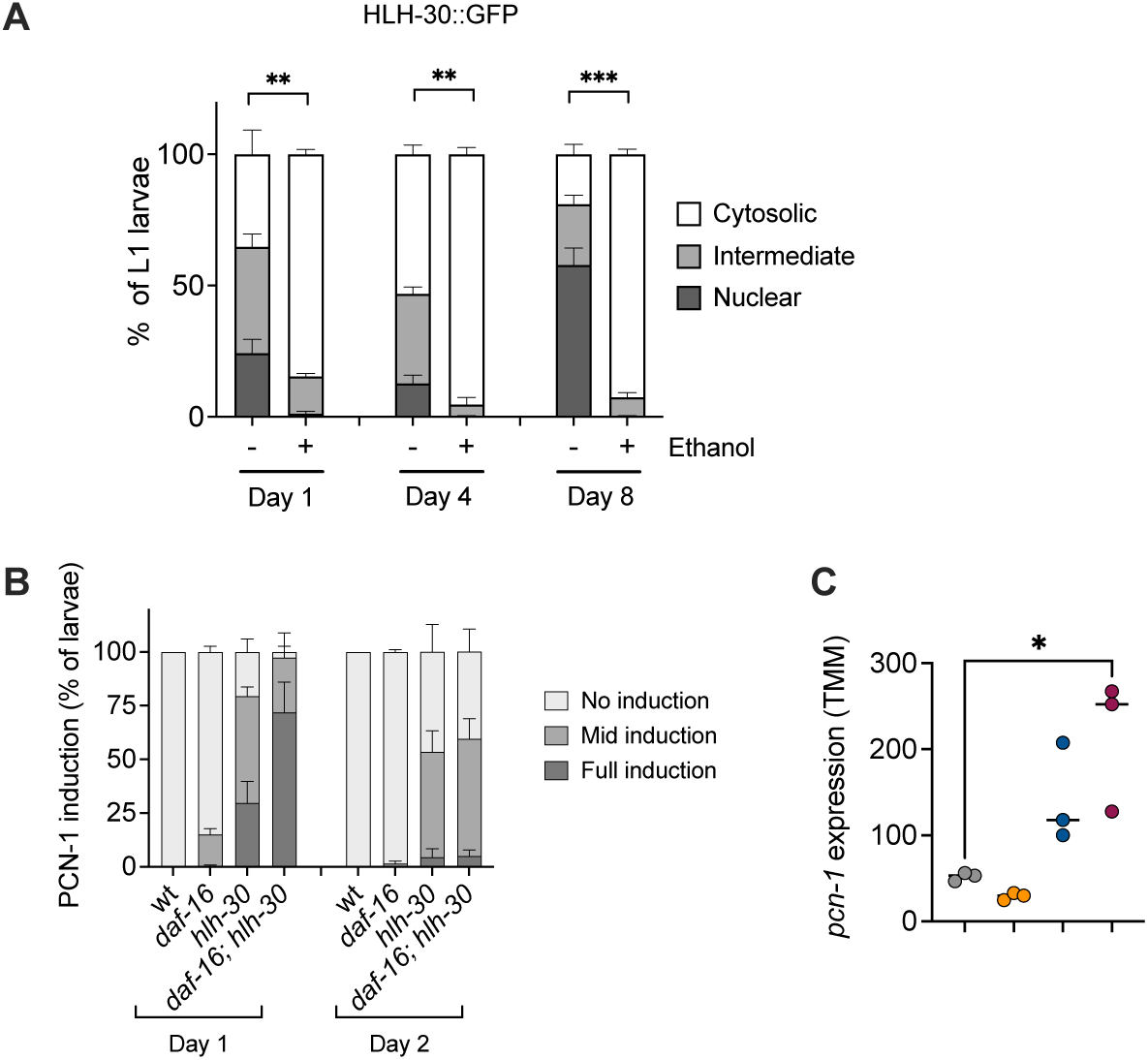
Relative to Figure 4. **(A)** Subcellular localization of HLH-30::GFP in M9 without (-) or with (+) 0.1 % (V/V) ethanol, after 1, 4 and 8 days of L1 starvation. Error bars represent SEM. We performed t-test to compare the effect of ethanol on the fraction of larvae with nuclear localization at each day of starvation. Only significant differences are shown. **(B)** Fraction of larvae with different levels of PCN-1 induction categorized as high, mid or no induction after 1 or 2 days of L1 starvation, for the wild type, *daf-16(mu86)*, *hlh-30(tm1978)*, and *daf-16;hlh-30*. **(C)** Expression levels of *pcn-1* after one of L1 starvation, for the wild type, *daf-16(mu86)*, *hlh- 30(tm1978)*, and *daf-16;hlh-30*. Results are shown as TMM (trimmed mean of M; File S2) values from the mRNA-seq experiment. For statistical analysis, we compared performed One-way ANOVA followed by Tukeýs multiple comparisons. Only significant differences to the wild-type strain are shown. Complete statistics report can be found in File S1.

**Figure S6.**
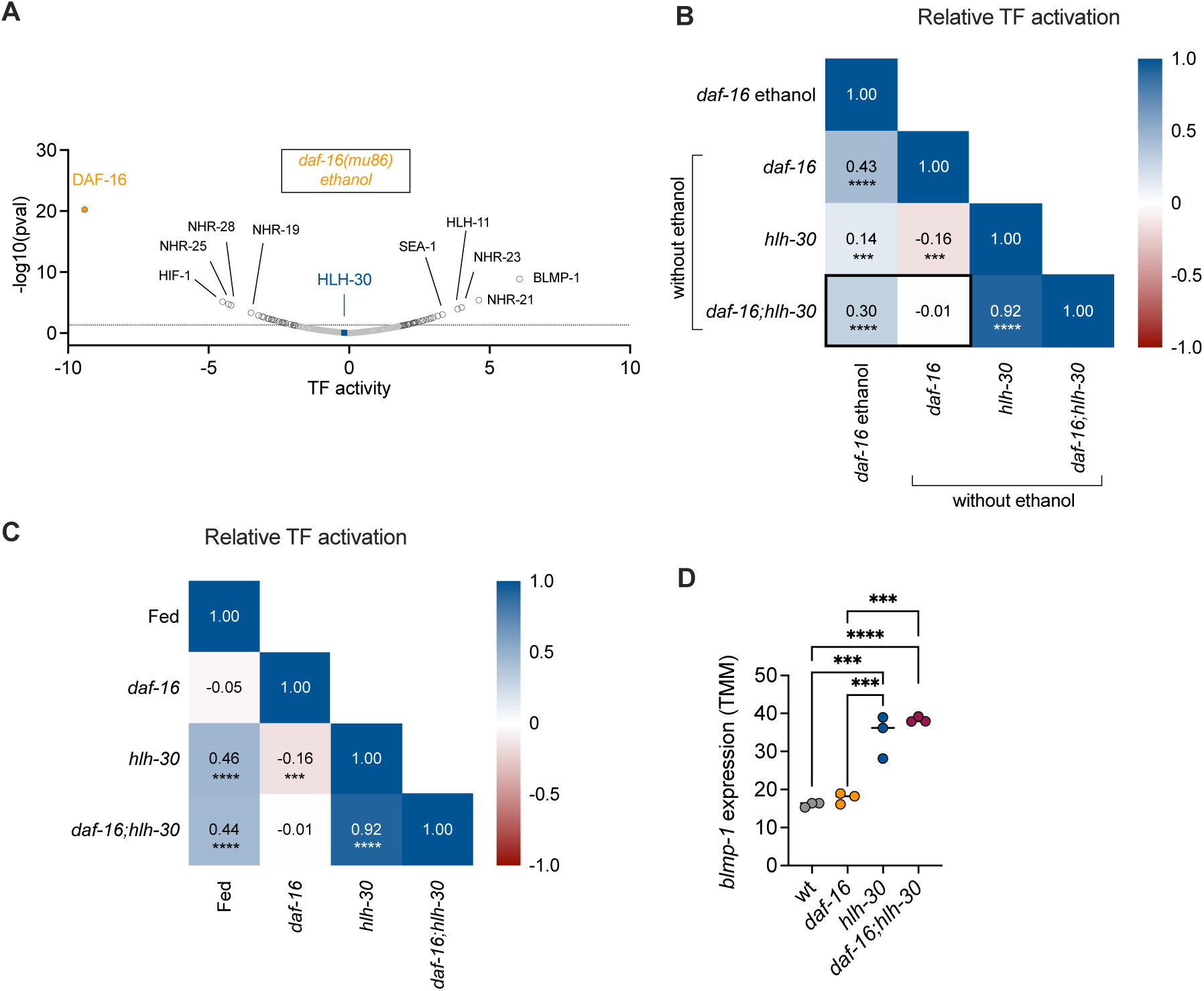
Relative to Figure 5. **(A)** Plot showing TF activity score for *daf-16* mutants versus wild type larvae, starved in the presence of ethanol. **(B)** Correlation matrix for TF activation scores of 487 transcription factors, for all possible pairs of conditions including *fed* (fed wt vs. starved wt), and *daf-16*, *hlh-30* and *daf-16;hlh-30* (vs. wt). **(C)** Correlation matrix for TF activation scores of 487 transcription factors, for all possible pairs of conditions including *daf-16* in ethanol (vs. wt in ethanol), and *daf-16*, *hlh-30* and *daf-16;hlh-30* without ethanol (vs. wt without ethanol). **(D)** Expression levels of *blmp-1* after one day of L1 starvation, for the wild type, *daf-16(mu86)*, *hlh-30(tm1978)*, and *daf-16;hlh-30*. Results are shown as TMM values from the mRNA-seq experiment. For statistical analysis, we compared performed One-way ANOVA followed by Tukeýs multiple comparisons. Complete statistics report can be found in File S1.

**Supplementary Table 1.**
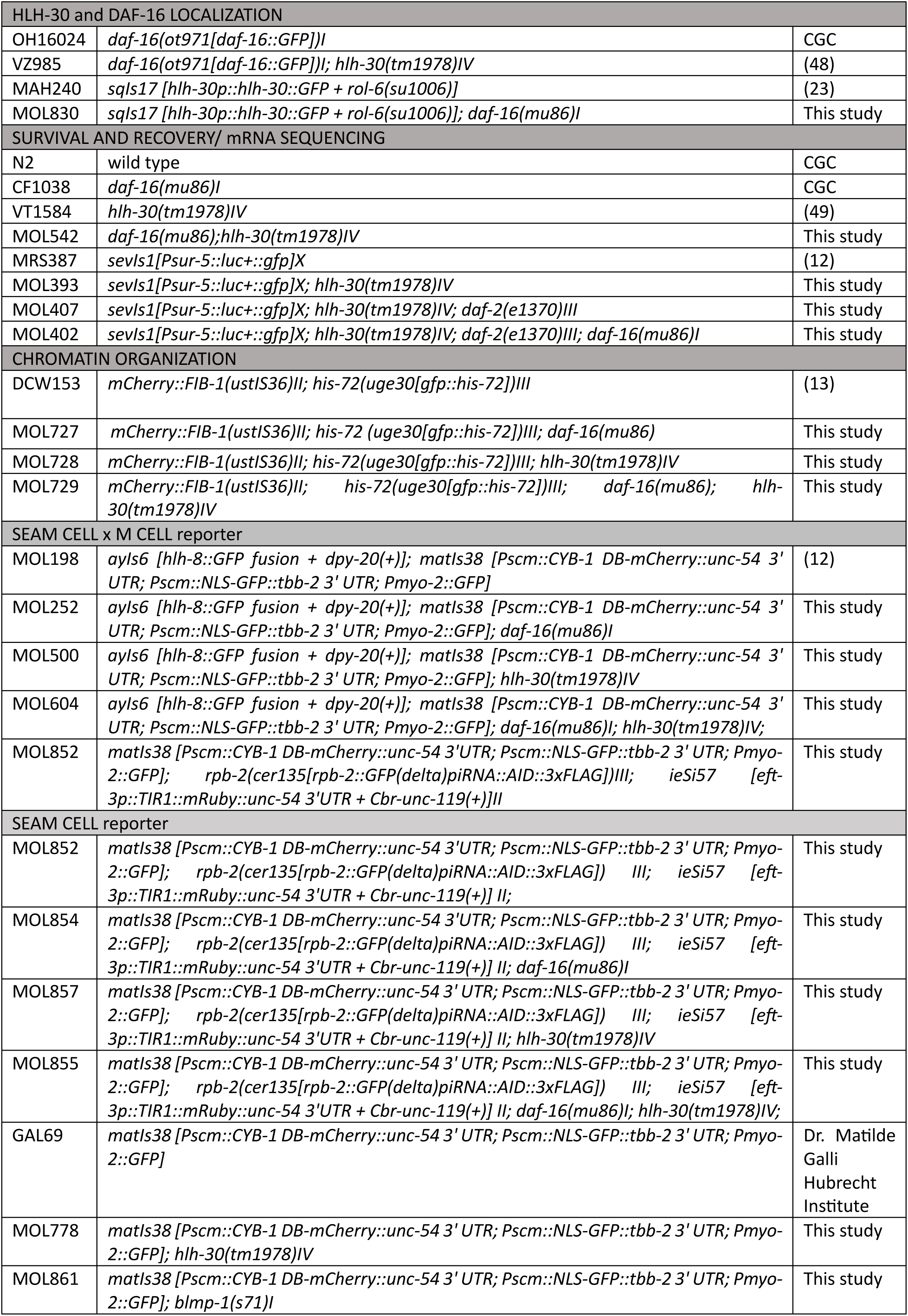
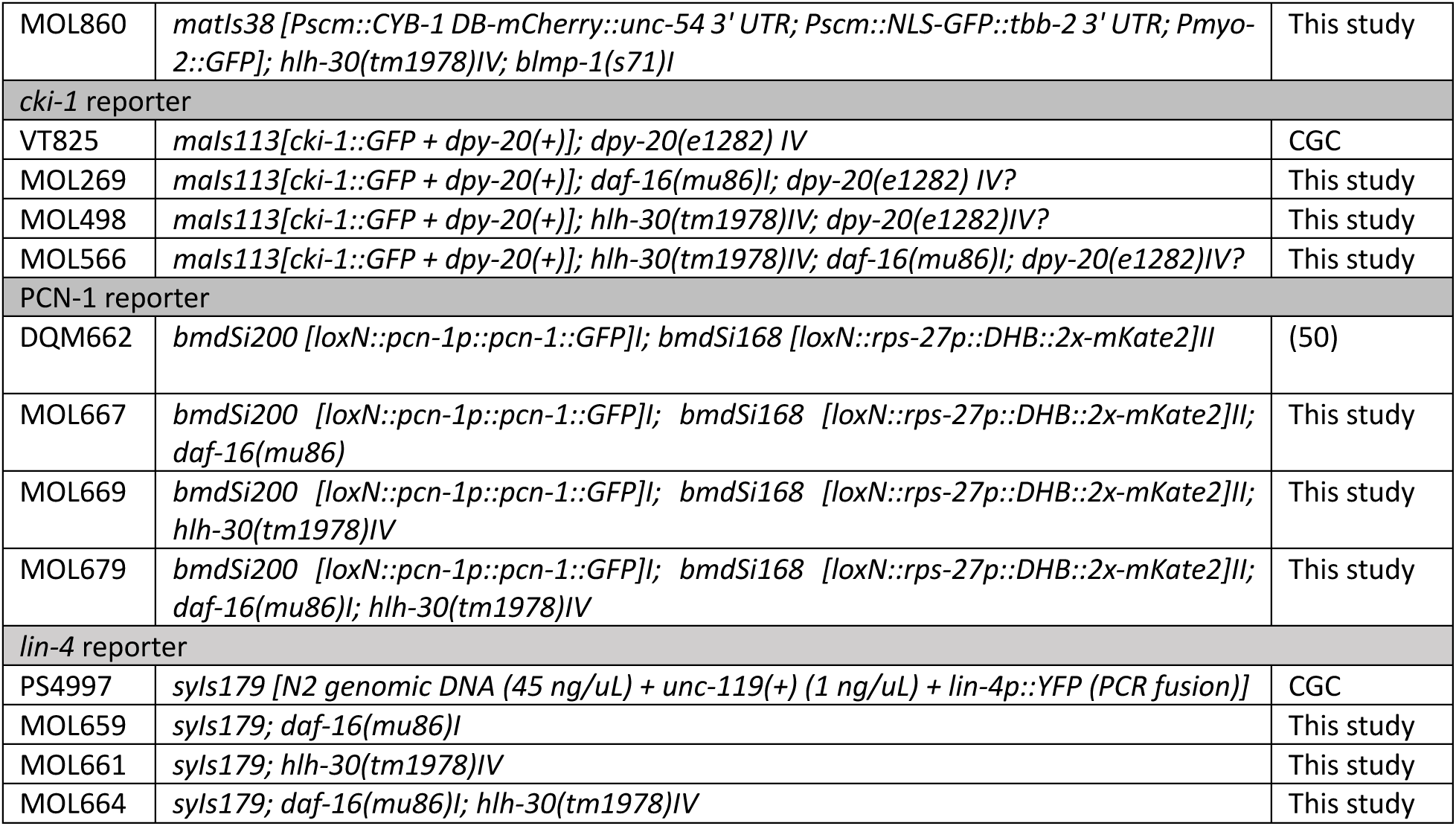
List of strains.

| HLH-30 and DAF-16 LOCALIZATION |  |  |
| --- | --- | --- |
| OH16024 | *daf-16(ot971[daf-16::GFP])I* | CGC |
| VZ985 | *daf-16(ot971[daf-16::GFP])I; hlh-30(tm1978)IV* | (48) |
| MAH240 | *sqls17 [hlh-30p::hlh-30::GFP + rol-6(su1006)]* | (23) |
| MOL830 | *sqls17 [hlh-30p::hlh-30::GFP + rol-6(su1006)]; daf-16(mu86)I* | This study |
| SURVIVAL AND RECOVERY/ mRNA SEQUENCING |  |  |
| N2 | wild type | CGC |
| CF1038 | *daf-16(mu86)I* | CGC |
| VT1584 | *hlh-30(tm1978)IV* | (49) |
| MOL542 | *daf-16(mu86); hlh-30(tm1978)IV* | This study |
| MRS387 | *sevls1[Psur-5::luc+::gfp]X* | (12) |
| MOL393 | *sevls1[Psur-5::luc+::gfp]X; hlh-30(tm1978)IV* | This study |
| MOL407 | *sevls1[Psur-5::luc+::gfp]X; hlh-30(tm1978)IV; daf-2(e1370)III* | This study |
| MOL402 | *sevls1[Psur-5::luc+::gfp]X; hlh-30(tm1978)IV; daf-2(e1370)III; daf-16(mu86)I* | This study |
| CHROMATIN ORGANIZATION |  |  |
| DCW153 | *mCherry::FIB-1(ustIS36)II; his-72(uge30[gfp::his-72])III* | (13) |
| MOL727 | *mCherry::FIB-1(ustIS36)II; his-72 (uge30[gfp::his-72])III; daf-16(mu86)* | This study |
| MOL728 | *mCherry::FIB-1(ustIS36)II; his-72(uge30[gfp::his-72])III; hlh-30(tm1978)IV* | This study |
| MOL729 | *mCherry::FIB-1(ustIS36)II; his-72(uge30[gfp::his-72])III; daf-16(mu86); hlh-30(tm1978)IV* | This study |
| SEAM CELL x M CELL reporter |  |  |
| MOL198 | *ayls6 [hlh-8::GFP fusion + dpy-20(+)]; matIs38 [Pscm::CYB-1 DB-mCherry::unc-54 3' UTR; Pscm::NLS-GFP::tbb-2 3' UTR; Pmyo-2::GFP]* | (12) |
| MOL252 | *ayls6 [hlh-8::GFP fusion + dpy-20(+)]; matIs38 [Pscm::CYB-1 DB-mCherry::unc-54 3' UTR; Pscm::NLS-GFP::tbb-2 3' UTR; Pmyo-2::GFP]; daf-16(mu86)I* | This study |
| MOL500 | *ayls6 [hlh-8::GFP fusion + dpy-20(+)]; matIs38 [Pscm::CYB-1 DB-mCherry::unc-54 3' UTR; Pscm::NLS-GFP::tbb-2 3' UTR; Pmyo-2::GFP]; hlh-30(tm1978)IV* | This study |
| MOL604 | *ayls6 [hlh-8::GFP fusion + dpy-20(+)]; matIs38 [Pscm::CYB-1 DB-mCherry::unc-54 3' UTR; Pscm::NLS-GFP::tbb-2 3' UTR; Pmyo-2::GFP]; daf-16(mu86)I; hlh-30(tm1978)IV;* | This study |
| MOL852 | *matIs38 [Pscm::CYB-1 DB-mCherry::unc-54 3' UTR; Pscm::NLS-GFP::tbb-2 3' UTR; Pmyo-2::GFP]; rpb-2(cer135[rpb-2::GFP(delta)piRNA::AID::3xFLAG])III; ieSi57 [eft-3p::TIR1::mRuby::unc-54 3' UTR + Cbr-unc-119(+)]II* | This study |
| SEAM CELL reporter |  |  |
| MOL852 | *matIs38 [Pscm::CYB-1 DB-mCherry::unc-54 3' UTR; Pscm::NLS-GFP::tbb-2 3' UTR; Pmyo-2::GFP]; rpb-2(cer135[rpb-2::GFP(delta)piRNA::AID::3xFLAG]) III; ieSi57 [eft-3p::TIR1::mRuby::unc-54 3' UTR + Cbr-unc-119(+)] II;* | This study |
| MOL854 | *matIs38 [Pscm::CYB-1 DB-mCherry::unc-54 3' UTR; Pscm::NLS-GFP::tbb-2 3' UTR; Pmyo-2::GFP]; rpb-2(cer135[rpb-2::GFP(delta)piRNA::AID::3xFLAG]) III; ieSi57 [eft-3p::TIR1::mRuby::unc-54 3' UTR + Cbr-unc-119(+)] II; daf-16(mu86)I* | This study |
| MOL857 | *matIs38 [Pscm::CYB-1 DB-mCherry::unc-54 3' UTR; Pscm::NLS-GFP::tbb-2 3' UTR; Pmyo-2::GFP]; rpb-2(cer135[rpb-2::GFP(delta)piRNA::AID::3xFLAG]) III; ieSi57 [eft-3p::TIR1::mRuby::unc-54 3' UTR + Cbr-unc-119(+)] II; hlh-30(tm1978)IV* | This study |
| MOL855 | *matIs38 [Pscm::CYB-1 DB-mCherry::unc-54 3' UTR; Pscm::NLS-GFP::tbb-2 3' UTR; Pmyo-2::GFP]; rpb-2(cer135[rpb-2::GFP(delta)piRNA::AID::3xFLAG]) III; ieSi57 [eft-3p::TIR1::mRuby::unc-54 3' UTR + Cbr-unc-119(+)] II; daf-16(mu86)I; hlh-30(tm1978)IV;* | This study |
| GAL69 | *matIs38 [Pscm::CYB-1 DB-mCherry::unc-54 3' UTR; Pscm::NLS-GFP::tbb-2 3' UTR; Pmyo-2::GFP]* | Dr. Matilde Galli Hubrecht Institute |
| MOL778 | *matIs38 [Pscm::CYB-1 DB-mCherry::unc-54 3' UTR; Pscm::NLS-GFP::tbb-2 3' UTR; Pmyo-2::GFP]; hlh-30(tm1978)IV* | This study |
| MOL861 | *matIs38 [Pscm::CYB-1 DB-mCherry::unc-54 3' UTR; Pscm::NLS-GFP::tbb-2 3' UTR; Pmyo-2::GFP]; blmp-1(s71)I* | This study |
| MOL860 | *matIs38 [Pscm::CYB-1 DB-mCherry::unc-54 3' UTR; Pscm::NLS-GFP::tbb-2 3' UTR; Pmyo-2::GFP]; hlh-30(tm1978)IV; blmp-1(s71)I* | This study |
| **cki-1 reporter** |  |  |
| VT825 | *mals113[cki-1::GFP + dpy-20(+)]; dpy-20(e1282) IV* | CGC |
| MOL269 | *mals113[cki-1::GFP + dpy-20(+)]; daf-16(mu86)I; dpy-20(e1282) IV?* | This study |
| MOL498 | *mals113[cki-1::GFP + dpy-20(+)]; hlh-30(tm1978)IV; dpy-20(e1282)IV?* | This study |
| MOL566 | *mals113[cki-1::GFP + dpy-20(+)]; hlh-30(tm1978)IV; daf-16(mu86)I; dpy-20(e1282)IV?* | This study |
| **PCN-1 reporter** |  |  |
| DQM662 | *bmdSi200 [loxN::pcn-1p::pcn-1::GFP]I; bmdSi168 [loxN::rps-27p::DHB::2x-mKate2]II* | (50) |
| MOL667 | *bmdSi200 [loxN::pcn-1p::pcn-1::GFP]I; bmdSi168 [loxN::rps-27p::DHB::2x-mKate2]II; daf-16(mu86)* | This study |
| MOL669 | *bmdSi200 [loxN::pcn-1p::pcn-1::GFP]I; bmdSi168 [loxN::rps-27p::DHB::2x-mKate2]II; hlh-30(tm1978)IV* | This study |
| MOL679 | *bmdSi200 [loxN::pcn-1p::pcn-1::GFP]I; bmdSi168 [loxN::rps-27p::DHB::2x-mKate2]II; daf-16(mu86)I; hlh-30(tm1978)IV* | This study |
| **lin-4 reporter** |  |  |
| PS4997 | *syIs179 [N2 genomic DNA (45 ng/uL) + unc-119(+)(1 ng/uL) + lin-4p::YFP (PCR fusion)]* | CGC |
| MOL659 | *syIs179; daf-16(mu86)I* | This study |
| MOL661 | *syIs179; hlh-30(tm1978)IV* | This study |
| MOL664 | *syIs179; daf-16(mu86)I; hlh-30(tm1978)IV* | This study |

**Supplementary Table 2.**
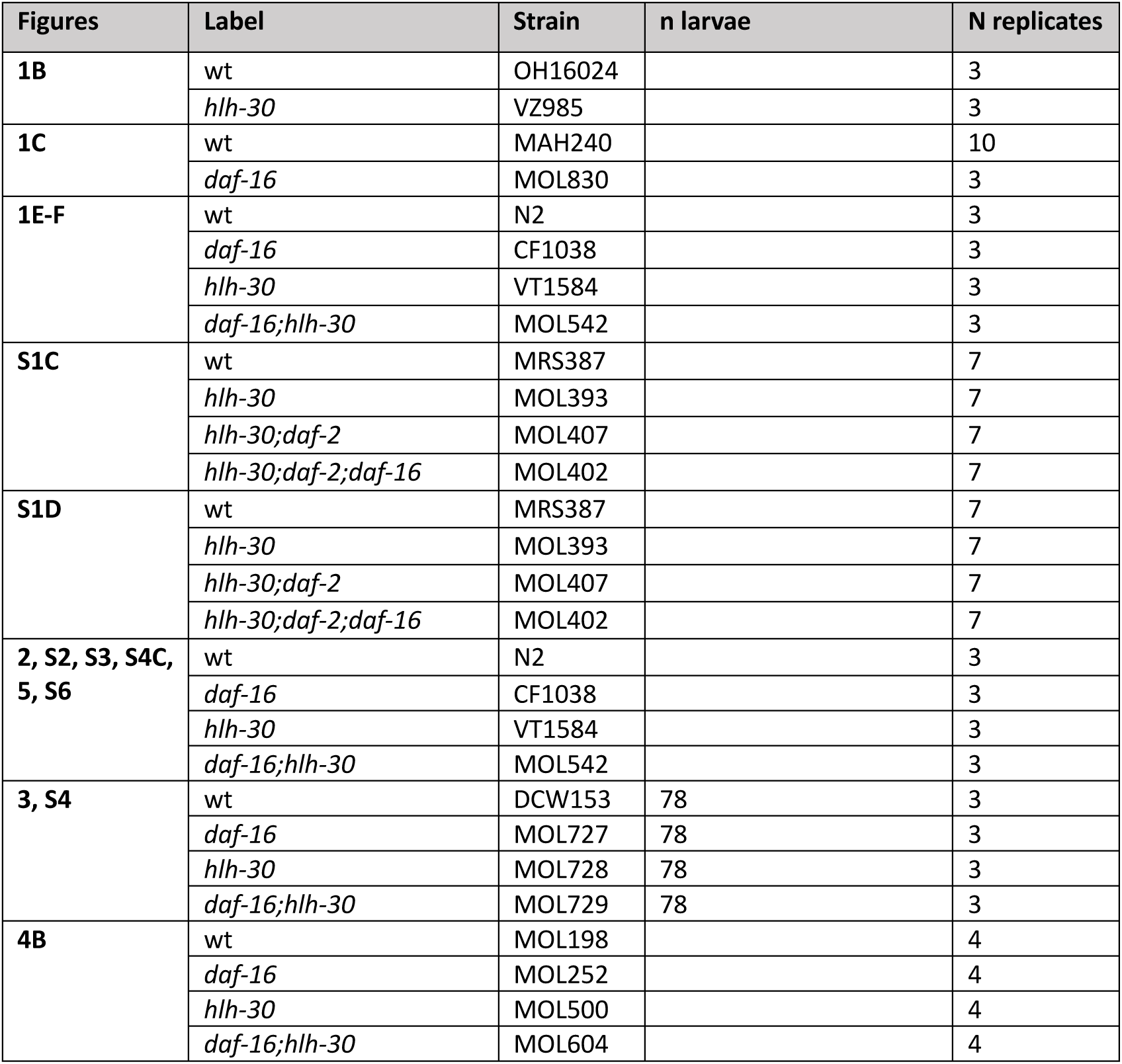
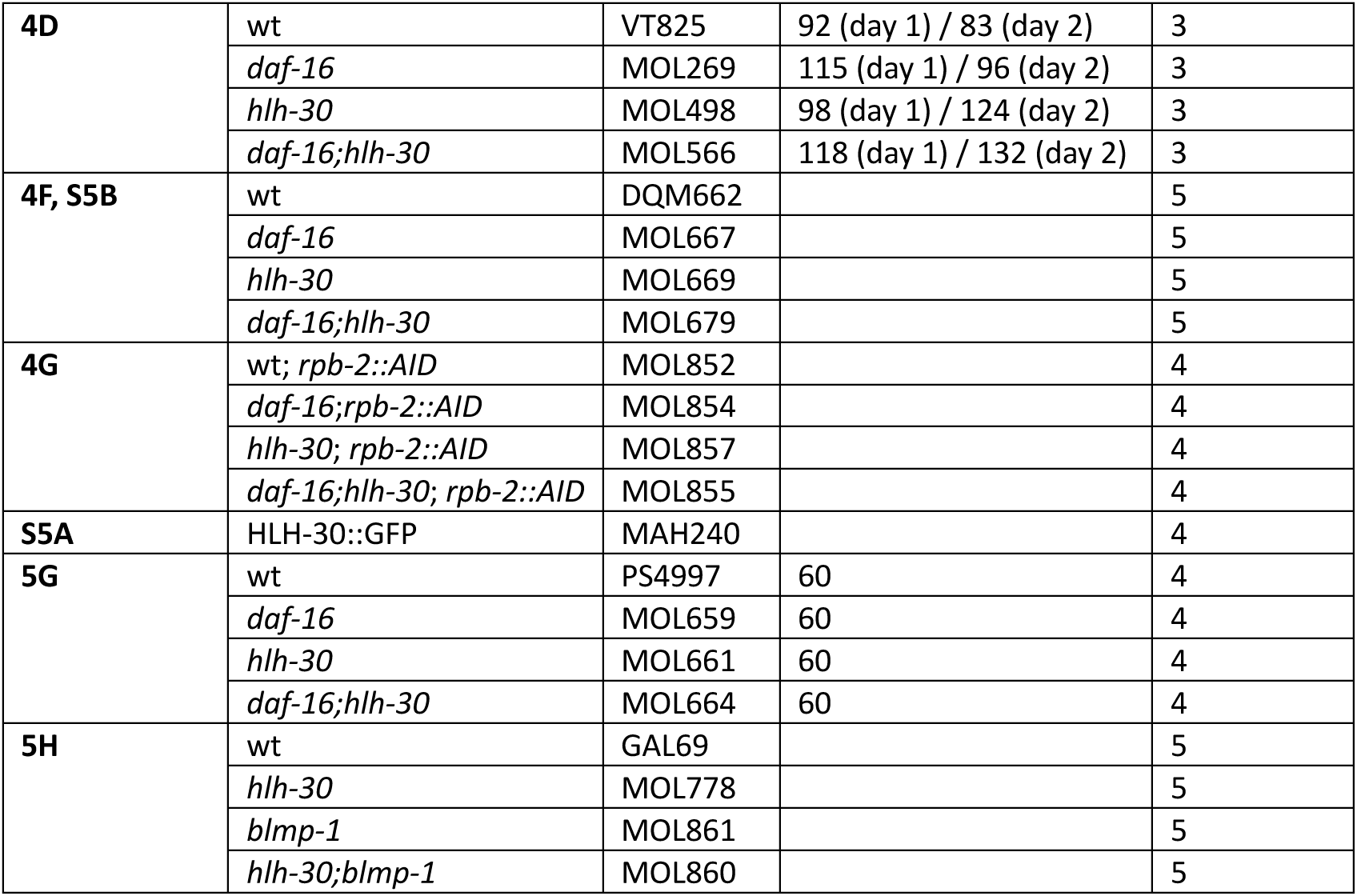
Strains, n larvae (when relevant) and N replicates per figure.

## REFERENCES

1. Murley, A., Wickham, K. & Dillin, A. Life in lockdown: Orchestrating endoplasmic reticulum and lysosome homeostasis for quiescent cells. Molecular Cell 82, 3526–3537 (2022).

2. Cheng, T. et al. Hematopoietic Stem Cell Quiescence Maintained by p21cip1/waf1. Science 287, 1804–1808 (2000).

3. Kwon, J. S. et al. Controlling Depth of Cellular Quiescence by an Rb- E2F Network Switch. CellReports 20, 3223–3235 (2017).

4. Fujimaki, K. et al. Graded regulation of cellular quiescence depth between proliferation and senescence by a lysosomal dimmer switch. Proceedings of the National Academy of Sciences 116, 22624–22634 (2019).

5. Oh, J., Lee, Y. D. & Wagers, A. J. Stem cell aging: mechanisms, regulators and therapeutic opportunities. Nat Med 20, 870–880 (2014).

6. Roux, A. E., Langhans, K., Huynh, W. & Kenyon, C. Reversible Age-Related Phenotypes Induced during Larval Quiescence in C. elegans. Cell Metabolism 23, 1113–1126 (2016).

7. Leeman, D. S. et al. Lysosome activation clears aggregates and enhances quiescent neural stem cell activation during aging. Science 359, 1277–1283 (2018).

8. Hidalgo San Jose, L., et al. Modest Declines in Proteome Quality Impair Hematopoietic Stem Cell Self-Renewal. CellReports 30, 69–80.e6 (2020).

9. Mata-Cabana, A., Romero-Expósito, F. J. & Olmedo, M. Aging during C. elegans L1 quiescence. Aging (Albany NY) 12, 17756–17758 (2020).

10. Murley, A. et al. Quiescent cell re-entry is limited by macroautophagy-induced lysosomal damage. Cell 188, 2670–2686.e14 (2025).

11. Hong, Y., Roy, R. & Ambros, V. Developmental regulation of a cyclin-dependent kinase inhibitor controls postembryonic cell cycle progression in *Caenorhabditis elegans*. Development 125, 3585–3597 (1998).

12. Olmedo, M. et al. Prolonged quiescence delays somatic stem cell-like divisions in Caenorhabditis elegans and is controlled by insulin signaling. Aging Cell 19, e13085 (2020).

13. Al-Refaie, N. et al. Fasting shapes chromatin architecture through an mTOR/RNA Pol I axis. Nat Cell Biol 26, 1903–1917 (2024).

14. Baugh, L. R. To Grow or Not to Grow: Nutritional Control of Development During Caenorhabditis elegans L1 Arrest. Genetics 194, 539–555 (2013).

15. Muñoz, M. J. & Riddle, D. L. Positive Selection of *Caenorhabditis elegans* Mutants With Increased Stress Resistance and Longevity. Genetics 163, 171–180 (2003).

16. Murphy, J. T. et al. Simple nutrients bypass the requirement for HLH-30 in coupling lysosomal nutrient sensing to survival. PLoS Biol. 17, e3000245–42 (2019).

17. Settembre, C. et al. TFEB controls cellular lipid metabolism through a starvation-induced autoregulatory loop. Nat Cell Biol 15, 647–658 (2013).

18. Lin, X.-X. et al. DAF-16/FOXO and HLH-30/TFEB function as combinatorial transcription factors to promote stress resistance and longevity. Nature Communications 1–15 (2018). doi:10.1038/s41467-018-06624-0

19. Weinkove, D., Halstead, J., Gems, D. & Divecha, N. Long-term starvation and ageing induce AGE-1/PI 3-kinase-dependent translocation of DAF-16/FOXO to the cytoplasm. BMC Biol 4, 1 (2006).

20. Mata-Cabana, A. et al. Social Chemical Communication Determines Recovery From L1 Arrest via DAF-16 Activation. Front Cell Dev Biol 8, 588686 (2020).

21. Webster, A. K., Chitrakar, R., Taylor, S. M. & Baugh, L. R. Alternative somatic and germline gene-regulatory strategies during starvation-induced developmental arrest. CellReports 41, 111473 (2022).

22. Wu, T., et al. clusterProfiler 4.0: A universal enrichment tool for interpreting omics data. Innovation (Camb) 2, 100141 (2021).

23. Lapierre, L. R. et al. The TFEB orthologue HLH-30 regulates autophagy and modulates longevity in Caenorhabditis elegans. Nature Communications 4, 2267 (2013).

24. Astanina, E., Bussolino, F. & Doronzo, G. Multifaceted activities of transcription factor EB in cancer onset and progression. Mol Oncol 15, 327–346 (2021).

25. Fukuyama, M., Kontani, K., Katada, T. & Rougvie, A. E. The C. elegans Hypodermis Couples Progenitor Cell Quiescence to the Dietary State. Curr. Biol. 25, 1241–1248 (2015).

26. Baugh, L. R. & Sternberg, P. W. DAF-16/FOXO Regulates Transcription of cki-1/Cip/Kip and Repression of lin-4 during C. elegans L1 Arrest. Current Biology 16, 780–785 (2006).

27. Oswal, N., Martin, O. M. F., Stroustrup, S., Bruckner, M. A. M. & Stroustrup, N. A hierarchical process model links behavioral aging and lifespan in C. elegans. PLoS Computational Biology 18, e1010415 (2022).

28. Perez, M. F. CelEst: a unified gene regulatory network for estimating transcription factor activities in C. elegans. Genetics 229, 255 (2025).

29. Chen, J., Chitrakar, R. & Baugh, L. R. DAF-18/PTEN protects LIN-35/Rb from CLP-1/CAPN- mediated cleavage to promote starvation resistance. Life Sci Alliance 8, e202403147 (2025).

30. Stec, N. et al. An Epigenetic Priming Mechanism Mediated by Nutrient Sensing Regulates Transcriptional Output during C.elegans Development. CURBIO 31, 809–826.e6 (2021).

31. Feinbaum, R. E. A. The Timing of lin-4 RNA Accumulation Controls the Timing of Postembryonic Developmental Events in Caenorhabditis elegans. 1–9 (1999).

32. Kinney, B. et al. A circadian-like gene network programs the timing and dosage of heterochronic miRNA transcription during C. elegans development. Developmental Cell 58, 2563–2579.e8 (2023).

33. Bulteau, R. & Francesconi, M. Real age prediction from the transcriptome with RAPToR. Nat. Methods 19, 969–975 (2022).

34. Ricaurte-Perez, C., Wall, P., Dubuisson, O., Bohnert, K. & Johnson, A. E. DAF-16/FOXO and HLH-30/TFEB comprise a cooperative regulatory axis controlling tubular lysosome induction in C. elegans. 1–27 (2024). doi:10.21203/rs.3.rs-4049366/v1

35. Castro, P. V., Khare, S., Young, B. D. & Clarke, S. G. Caenorhabditis elegans Battling Starvation Stress: Low Levels of Ethanol Prolong Lifespan in L1 Larvae. PLoS ONE 7, e29984 (2012).

36. Bar-Peled, L. & Sabatini, D. M. Regulation of mTORC1 by amino acids. Trends in Cell Biology 24, 400–406 (2014).

37. Horn, M. et al. DRE-1/FBXO11-Dependent Degradation of BLMP-1/BLIMP-1 Governs C. elegans Developmental Timing and Maturation. Developmental Cell 28, 697–710 (2014).

38. Hauser, Y. P., Meeuse, M. W. M., Gaidatzis, D. & Großhans, H. The BLMP-1 transcription factor promotes oscillatory gene expression to achieve timely molting. bioRxiv 9, 117– 43 (2022).

39. Bikoff, E. K., Morgan, M. A. & Robertson, E. J. An expanding job description for Blimp- 1/PRDM1. Curr. Opin. Genet. Dev. 19, 379–385 (2009).

40. Napolitano, G. & Ballabio, A. TFEB at a glance. J Cell Sci 129, 2475–2481 (2016).

41. Doronzo, G. et al. TFEB controls vascular development by regulating the proliferation of endothelial cells. The EMBO Journal 38, (2019).

42. Pisonero-Vaquero, S., Soldati, C., Cesana, M., Ballabio, A. & Medina, D. L. TFEB Modulates p21/WAF1/CIP1 during the DNA Damage Response. Cells 9, (2020).

43. Love, M. I., Huber, W. & Anders, S. Moderated estimation of fold change and dispersion for RNA-seq data with DESeq2. Genome Biol. 15, 550 (2014).

44. Wickham, H. ggplot2. 1–266 (2016).

45. Gu, Z., Eils, R. & Schlesner, M. Complex heatmaps reveal patterns and correlations in multidimensional genomic data. Bioinformatics 32, 2847–2849 (2016).

46. Padovani, F., Mairhörmann, B., Falter-Braun, P., Lengefeld, J. & Schmoller, K. M. Segmentation, tracking and cell cycle analysis of live-cell imaging data with Cell-ACDC. BMC Biol 20, 174–18 (2022).

47. Lord, S. J., Velle, K. B., Mullins, R. D. & Fritz-Laylin, L. K. SuperPlots: Communicating reproducibility and variability in cell biology. The Journal of Cell Biology 219, (2020).

48. Colino-Lage, H. et al. Regulation of *Caenorhabditis elegans* HLH-30 subcellular localization dynamics: Evidence for a redox-dependent mechanism. Free Radic Biol Med 223, 369–383 (2024).

49. Grove, C. A. et al. A Multiparameter Network Reveals Extensive Divergence between *C. elegans* bHLH Transcription Factors. Cell 138, 314–327 (2009).

50. Adikes, R. C. et al. Visualizing the metazoan proliferation-quiescence decision in vivo. eLife 9:e63265 (2020).

